# The spinal premotor network driving scratching flexor and extensor alternation

**DOI:** 10.1101/2025.01.08.631866

**Authors:** Mingchen Yao, Akira Nagamori, Eiman Azim, Tatyana Sharpee, Martyn Goulding, David Golomb, Graziana Gatto

**Author notes:** Co-corresponding authors Corresponding author contacts: Eiman Azim, Salk Institute for Biological Studies, Tatyana Sharpee, Salk Institute for Biological Studies, Martyn Goulding, Salk Institute for Biological Studies, David Golomb, Ben Gurion University, Graziana Gatto, University Hospital of Cologne.

## Abstract

Rhythmic motor behaviors are generated by neural networks termed central pattern generators (CPGs). Although locomotor CPGs have been extensively characterized, it remains unknown how the neuronal populations composing them interact to generate adaptive rhythms. We explored the non-linear cooperation dynamics among the three main populations of ipsilaterally projecting spinal CPG neurons – V1, V2a, V2b neurons – in scratch reflex rhythmogenesis. Ablation of all three neuronal subtypes reduced the oscillation frequency. Activation of excitatory V2a neurons enhanced the oscillation frequency, while activating inhibitory V1 neurons caused atonia. These findings required the development of a novel neuromechanical model that consists of flexor and extensor modules coupled via inhibition, in which rhythm in each module is generated by self-bursting excitatory populations and accelerated by intra-module inhibition. Inter-module inhibition coordinates the phases of flexor and extensor activity and slows the oscillations, while facilitation mechanisms in excitatory neurons explain the V2a activation-driven increase in frequency.

## Introduction

Diverse rhythmic motor behaviors occur across the animal kingdom, arguing evolution has selected several mechanisms to generate rhythmic activity. However, the neuron types and circuit dynamics underlying this diversity of rhythms remain a long-standing question. Conserved neuronal networks in the vertebrate spinal cord, termed central pattern generators (CPGs) are able to produce a range of rhythmic motor behaviors including swimming, walking, running and scratching. These rhythmic behaviors show specificity in the timing and coordination patterns of muscle contractions, largely by shaping the output of the different motor pools that innervate limb and appendicular muscles (Goulding, 2009; Kiehn, 2016).

Locomotion has long been used as model behavior to study the mechanistic underpinnings of rhythm generation and the underlying cellular and circuit components of CPGs in aquatic and terrestrial vertebrates. The spinal locomotor CPGs comprise six genetically identified interneuron populations - dI6, V0, V1, V2a, V2b and V3 neurons - that differ in their axonal projection patterns and neurotransmitter phenotypes (Goulding, 2009). Studies in mammals have revealed that the locomotor rhythm is primarily generated by excitatory neurons and/or neural networks that display intrinsic bursting properties (Dougherty and Ha, 2019). These neurons are primarily ipsilaterally projecting such that each spinal hemicord has the capacity to generate rhythmic activity (Kiehn, 2016). In zebrafish, the excitatory Chx10-expressing V2a neurons (Chx10-V2a) have been shown to be rhythmogenic (Ampatzis et al., 2014). However, in rodents ablating or silencing Chx10-V2a neurons does not abolish the locomotor rhythm (Crone et al., 2008, 2009; Dougherty et al., 2013). This may be due to compensatory activity from Shox2-expressing V2a neurons (Shox2-V2a) (Dougherty et al., 2013), V3 neurons (Zhang et al., 2008), and the non-canonical CPG population, the Hb9-expressing neurons (Caldeira et al., 2017). Remarkably, inhibitory neurons also contribute to rhythmogenesis, with the loss of V1 neurons leading to a slowing of the fictive locomotor rhythm (Gosgnach et al., 2006; Trevisan et al., 2024). Fast swimming in zebrafish (Kimura and Higashijima, 2019) and tadpoles (Vijatovic et al., 2024) also depends on V1 neuron activity. While the loss of the ipsilateral inhibitory V2b neurons increases swimming frequency in larval zebrafish (Callahan et al., 2019), ablating these neurons in adult mice was found to have little or no effect on the locomotor rhythm (Britz et al., 2015). Despite these advances in defining the functional contribution of V1, V2a and V2b neurons to locomotor rhythmogenesis, we are still far from understanding how the dynamic interactions among these excitatory and inhibitory neurons contribute to generate other rhythmic motor behaviors.

Current models of how excitatory and inhibitory neurons cooperate to drive rhythm and motoneuron firing (Brown, 1911, 1914; Friesen and Stent, 1977; McCrea and Rybak, 2008; Perkel and Mulloney, 1974) have coalesced around the classic half-center model driven by the inhibitory coupling of the flexor and extensor modules (Brown, 1911). In alternative models, locomotor rhythm arises from individual modules acting as intrinsic bursting oscillators (Ausborn et al., 2018; Dougherty and Ha, 2019; Jankowska et al., 1967a, 1967b; Lundberg, 1981; Wallén and Grillner, 1987). Nevertheless, it remains unclear if these rhythmogenic mechanisms and their underlying neural components and circuit architectures hold true for other rhythmic behaviors, given that different motor behaviors such as swimming and walking in aquatic vertebrates rely on both shared and specialized CPG neurons (Berkowitz et al., 2010). Molecularly distinct subsets of interneurons and motoneurons that are organized into specialized microcircuit modules enable adult zebrafish to swim at different speeds (Ampatzis et al., 2014). It is not known, however, whether the same principles apply to rhythmic movements in limbed animals. The mechanisms driving faster rhythmic movements in limbed animals have been particularly difficult to dissect, as animals adapt to higher speeds not only by simply increasing frequency but also by changing gait (Bellardita and Kiehn, 2015; Crone et al., 2008; Pinto and Golubitsky, 2006; Talpalar et al., 2013; Zhang et al., 2022). To address these questions we focused on a simpler rhythmic behavior, the scratch reflex (Sherrington, 1910). Animals typically walk at a frequency of 2-3 Hz (Bellardita and Kiehn, 2015), but scratch at a frequency of 6 Hz (Frigon, 2012), which here we refer to as a ‘fast rhythm’. A further advantage is that the scratch rhythm is unilateral and mostly mono-articular, thus allowing us to focus on the ipsilaterally projecting excitatory V2a and inhibitory V1 and V2b neurons to define the circuit architecture and the neural dynamics that underlie ‘fast rhythm’ generation in limbed animals.

We found that ablation of either excitatory V2a or inhibitory V1 and V2b neurons reduced the frequency of scratch oscillations. Activation of V2a neurons increased the cadence. By contrast, activating V1 neurons caused an atonia-like state. Our experimental results contrasted previous models predictions, which for example suggest that reducing inhibition should result in faster rhythmic oscillations (*e*.*g*. (Ausborn et al., 2021; Rybak et al., 2006; Sherwood et al., 2011; Skinner et al., 1994)). These incongruities prompted us to develop a novel neuromechanical model, in which flexor and extensor modules are coupled via mutual inhibition. The rhythm in our model is generated by self-bursting excitatory populations within each module and is dependent on the relative strength of intra- and inter-module inhibition. Facilitation mechanisms in excitatory neurons explain the faster scratch oscillations caused by activation of V2a neurons. In sum, the computational modeling based on our experimental manipulations provides a mechanistic solution to the non-linear cooperation dynamics that drive fast rhythmic behaviors in an ipsilateral network of excitatory and inhibitory neurons.

## Results

### High-frequency flexor and extensor oscillations during scratching

Scratching is an evolutionary conserved reflex, executed as a stereotypical sequence of high frequency hindlimb oscillations (Sherrington, 1910). In a series of preliminary experiments, we analyzed the kinematic parameters of scratching in mice by injecting chloroquine, a potent pruritogen, in the nape of mice (**Figure 1A**). Scratch behavior was recorded for 30 minutes (**Video S1**), showing a sustained response with multiple scratch episodes of variable duration and frequency (**Figure 1B**). Individual scratch episodes were comprised of three phases: approach, rhythmic oscillations, and termination (**Figure 1C, 1D**), in which the hindlimb is initially flexed towards the trunk (approach), then rhythmically extended and flexed around the nape (oscillations), and finally extended back towards the floor (termination) (**Figure 1D**). Kinematic tracking showed that mice predominantly use the distal limb to execute the movement, with scratching generated primarily by rhythmic oscillations of the ankle (**Figure 1D, 1E**). The approach phase involved a stereotypical flexion of the limb with a similar duration across episodes and animals (**Figure S1C**), however, the frequency of oscillations, the duration of individual episodes, the amplitude of movements, and the termination phase were found to be highly variable within and across trials and mice (**Figure S1A, S1B, S1D, S1E**). By quantifying the frequency distribution of scratch oscillations, we were able to evaluate the intrinsic motor variability of this behavior that displayed an oscillation frequency ranging from 2 to 15 Hz, with a peak at 6 Hz (**Figure 1F**). These analyses demonstrate that mouse scratching is a unilateral movement driven by ankle joint oscillations over a range of frequencies, and it confirmed its use as an experimentally accessible model to address how CPG neurons generate and modulate motor rhythm.

**Figure 1.**
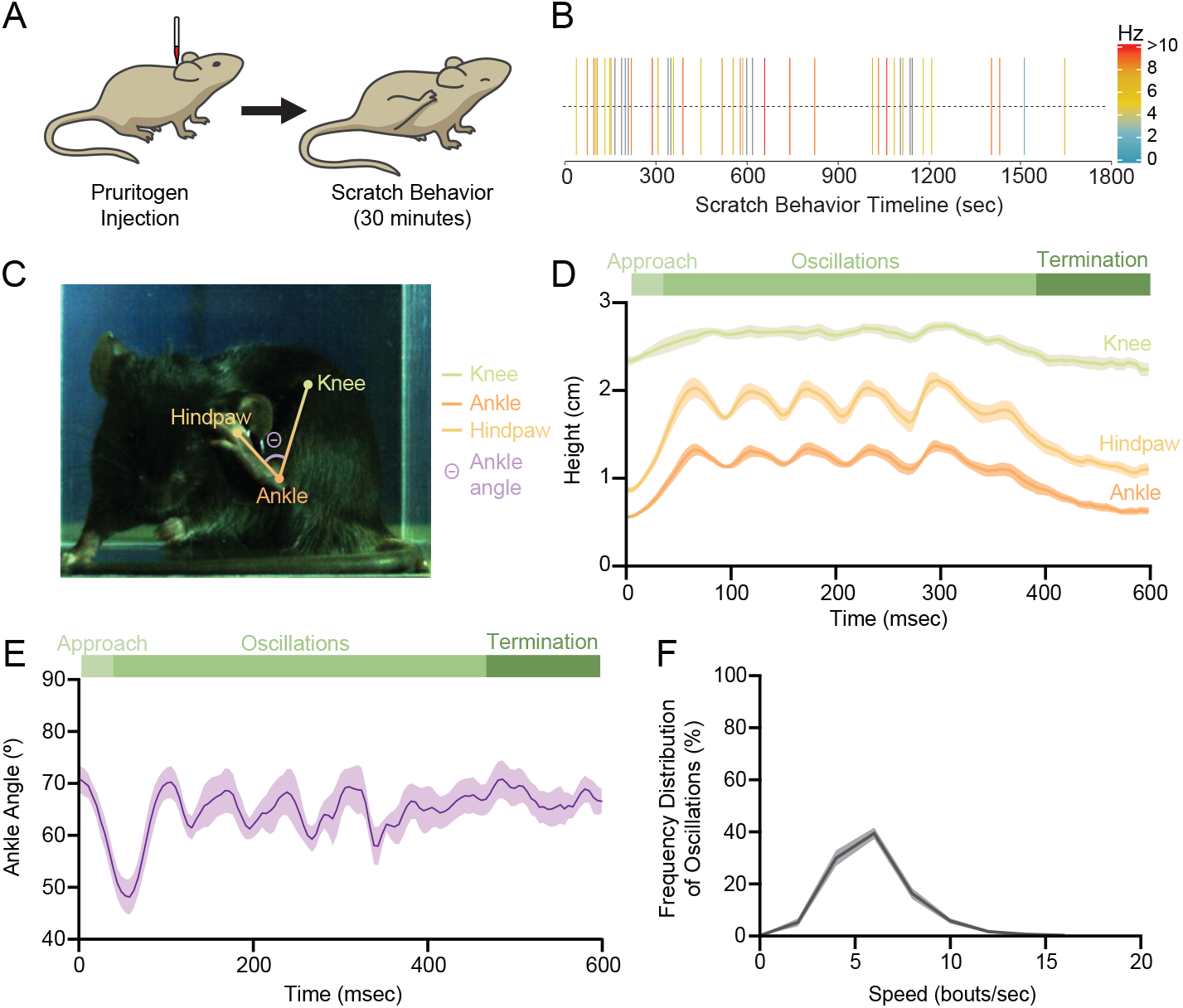
High-frequency flexor and extensor oscillations during scratching. A) Schematic illustrating the injection of a pruritogen (chloroquine) in the nape of mice to induce scratching. B) Raster plot showing multiple episodes of scratching in a single wild-type mouse following the injection of chloroquine. Frequency of oscillations of each episode is color-coded, as indicated in the legend. C) Image of a mouse during the oscillation phase of scratching, with tracked knee (green), ankle (dark orange), and hindpaw (light orange) landmarks. The ankle angle (θ) is indicated in purple. D,E) Line graphs representing the height (y-coordinates) of knee (green), ankle (dark orange) and hindpaw (light orange) (D) and the ankle angle (E) during scratching, N=10 episodes from 3 mice. Episodes with similar duration and frequency across three animals were chosen and averaged. F) Frequency distributions of the speed of oscillations (bouts/second) of all episodes occurring in the 30 minutes of recorded scratch response, N=30 mice. A bout is defined as the completion of one circular trajectory around the nape (one oscillation). Data presented as mean ± SEM, SEM represented as shaded area.

### Ipsilateral excitatory V2a neurons drive the high-frequency scratch oscillations

Rhythmic movements are driven by CPG neurons residing in the ventral spinal cord (**Figure 2A**) (Goulding, 2009). Since scratching is a hindlimb unilateral rhythmic movement, we focused on the three CPG neuron populations with ipsilateral projections: the excitatory V2a neurons and the inhibitory V1 and V2b neurons. The activity of these neuronal populations was selectively manipulated using an intersectional genetic approach involving mouse crosses that combined the *hCdx2::FlpO* allele (Britz et al., 2015), which restricts expression to the spinal cord, with selected *Cre* lines that developmentally capture the three neuronal lineages (**Figure 2B**). Dual recombinase-expressing mice were then crossed with animals carrying one of among these *Cre-* and *Flp*-dependent effectors, the diphtheria toxin receptor (DTR) allele (*Tau*^*ds-DTR*^) (Britz et al., 2015) for neuronal ablation, the hM4Di DREADD (*R26*^*ds-hM*4*Di*^) (Bourane et al., 2015) for transient neuronal silencing, and the hM3Dq DREADD (*R26*^*ds-hM*3*Dq*^) (Sciolino et al., 2016) for transient neuronal activation. Importantly, these receptors only become active following the administration of their ligands, thus allowing for the normal development of the spinal circuits.

**Figure 2.**
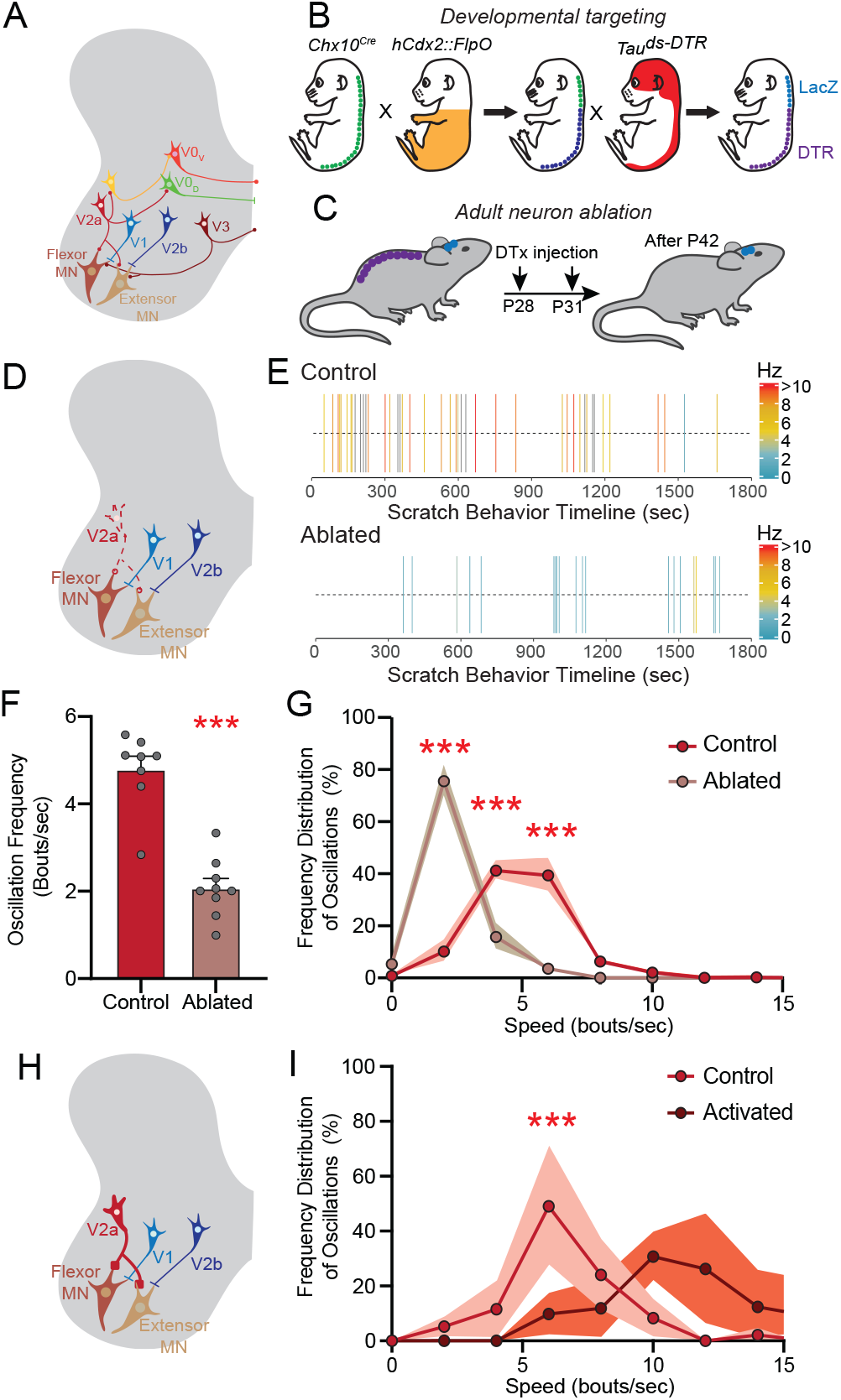
Ipsilateral excitatory V2a neurons drive high-frequency scratch oscillations. A-B,D,H) Schematics illustrating the neuronal populations in the ventral spinal cord constituting the central pattern generators (CPGs) [MN: motoneurons], the intersectional genetic approach to drive the expression of DTR (*Tau*^ds-DTR^) in spinal excitatory V2a neurons using *hCdx2::FlpO* and *Chx10*^Cre^ (B), the timeline of DTx injection and behavioral testing (C), and the ablation (D) and the CNO-driven activation (H) of V2a neurons. E) Raster plots showing the occurrence of scratching episodes in control and V2a neuron-ablated mice following injection of chloroquine. Frequency of oscillations of each episode is color-coded, as indicated in the legend. F) Bar graph showing the reduction in average oscillation frequency in V2a neuron-ablated mice compared to controls. Statistical analysis performed using two-tailed Student’s *t-test, p < 0*.*0001*. G) Frequency distributions of the speed of oscillations (bouts/second) of all episodes occurring in the 30 minutes of recorded scratch responses in control (red, N=8) and V2a neuron-ablated (brown, N=9) mice. Statistical analysis performed using two-way ANOVA (interaction genotype x speed, *p < 0*.*0001*) with Bonferroni’s *post-hoc* test, *p < 0*.*0001* for genotype comparison at 2, 4, and 6 bouts/sec. I) Frequency distributions of oscillations (bouts/second) during spontaneous scratching in controls (red, N=4) and in mice with CNO-activation of V2a neurons (dark red, N=7). Statistical analysis performed using two-way ANOVA (interaction genotype x speed, *p = 0*.*0004*) with Bonferroni’s *post-hoc* test, *p < 0*.*0001* for genotype comparison at 6 bouts/sec. Data presented as mean ± SEM. Individual mice represented as filled grey circles in F, SEM as shaded area in G and I.

Excitatory V2a neurons were ablated using *Chx10*^*Cre*^ and *hCdx2::FlpO* to drive the expression of DTR (Britz et al., 2015) and administering the diphtheria toxin (DTx) twice in adult animals (**Figure 2B, 2C**). Littermates lacking the *FlpO* allele were used as controls. Two weeks after injecting DTx we tested the motor dynamics during chloroquine-induced scratching (**Figure 2C**). Ablating V2a neurons (**Figure 2D, S2A**) strongly affected the scratch response (**Figure 2E**) by significantly reducing the total number of scratch bouts (**Figure S2B**) and the average frequency of scratching oscillations (**Figure 2F**). A frequency distribution analysis of all the scratch episodes showed a marked shift towards slow oscillations, with the frequency peaking at 2 Hz in V2a neuron-ablated mice (**Figure 2G**). To confirm that these results were not due to either the re-organization of the CPG circuitry post-ablation or compensation, we performed acute silencing experiments using the inhibitory DREADD hM4Di. CNO-dependent silencing of the V2a neurons (**Figure S2C**) also slowed the scratch rhythm (**Figure S2D-S2F**), phenocopying the impairments seen in scratching movements following V2a neuron ablation (**Figure 2F, 2G**).

We then activated spinal V2a neurons using our intersectional genetic approach to drive the expression of the excitatory DREADD hM3Dq in these neurons (**Figure 2H, S2G**). Mice in which the V2a neurons were activated with CNO generated similar number of spontaneous scratch bouts (**Figure S2H**) but faster scratch oscillations as compared to CNO-treated control littermates (**Figure 2I, S2I**). Taken together, our data show that V2a neuron ablation or silencing slows the scratch oscillation frequency, whereas V2a neuron activation increases it.

### Ipsilateral inhibitory neurons modulate the frequency of scratch oscillations

To assess the role of inhibitory populations in driving scratch oscillations we targeted the V1 and V2b neurons that are the principal source of ipsilateral inhibition onto limb motoneurons (Zhang et al., 2014). Expression of DTR was restricted to V1 neurons using *hCdx2::FlpO* and *En1*^*Cre*^ (Britz et al., 2015), with littermates lacking the *FlpO* allele used as controls. Ablating the V1 neurons (**Figure 3A, S3A**) markedly impaired the execution of scratching (**Figure 3B**) by decreasing the oscillation frequency (**Figure 3C, 3D**) and reducing the total number of scratch bouts (**Figure S3B**). Acute silencing of V1 neurons via CNO-driven stimulation of hM4Di (**Figure S3C**) recapitulated the ablation phenotype (**Figure S3D-S3F**), again confirming that the slowing of the rhythm was not due to ablation-induced re-wiring or compensatory mechanisms. In contrast, CNO-activation of V1 neurons via hM3Dq (**Figure 3E**) did not increase the speed of scratching, but resulted in an atonia-like state, with no movements observed until the drug wash-out (**Figure 3F**).

**Figure 3.**
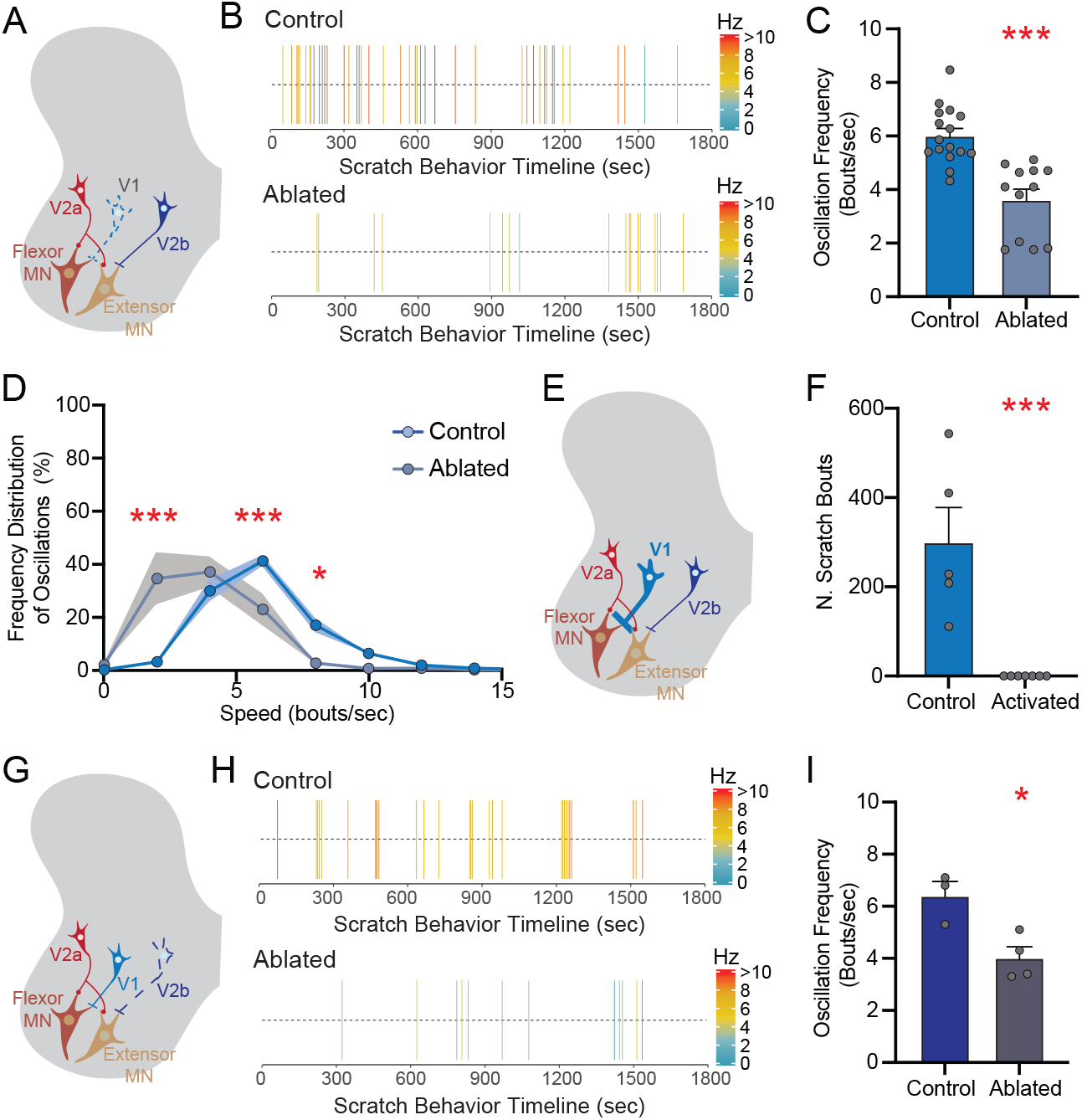
Ipsilateral inhibitory neurons modulate the frequency of scratch oscillations. A,E,G) Schematics illustrating the ablation (A) and the CNO-driven activation (E) of V1 neurons, and the ablation of V2b neurons (G). B) Raster plots showing the occurrence of scratching episodes in control and V1 neuron-ablated mice following injection of chloroquine. C) Bar graph showing the reduction in average oscillation frequency in V1 neuron-ablated mice compared to controls, *p < 0*.*0001*. D) Frequency distributions of the speed of oscillations (bouts/second) of all episodes occurring in the 30 minutes of recorded scratch response in control (azure, N=16) and V1 neuron-ablated (cerulean, N=12) mice. SEM represented as shaded area. Statistical analysis performed using two-way ANOVA (interaction genotype x speed, *p < 0*.*0001*) with Bonferroni’s *post-hoc* test, genotype comparison *p < 0*.*0001* at 2 bouts/sec, *p = 0*.*0004*, at 6 bouts/sec, *p = 0*.*0105* at 8 bouts/sec. F) Bar graph showing the lack of motor response (atonia state) in mice following CNO-induced activation of V1 neurons, *p = 0*.*0009*. H) Raster plots showing the occurrence of scratching episodes in control and V2b neuron-ablated mice following the injection of chloroquine. I) Bar graph showing the reduction in scratching oscillation frequency in V2b neuron-ablated mice compared to controls, *p = 0*.*0178*. Data presented as mean±SEM. Individual mice represented as filled grey circles. In raster plots in panels B and H the frequency of each episode is color-coded, as indicated in the legend. Statistical analysis performed using two-tailed Student’s *t-test*, unless otherwise indicated.

Due to the lack of a specific Cre line to target V2b neurons, we used *Hes2*^*iCre*^ to label all V2 neurons (Hayashi et al., 2023) combined with *VGAT*^*FlpO*^ to restrict DTR expression to the inhibitory V2b subset. DTx was injected intrathecally to ensure spinal specificity. Ablation of the V2b neurons (**Figure 3G, S3G**) significantly reduced the frequency of scratch oscillations (**Figure 3H, 3I, S3I**) compared to controls, with no changes to the total number of scratch bouts (**Figure S3H**). Taken together, our findings show that the inhibitory V1 and V2b neuron populations play a critical role in sustaining the high-frequency rhythmic oscillation during scratching. Furthermore, activation of the V1 neurons is sufficient to induce atonia, preventing the spontaneous execution of the scratch reflex.

### A neural model for high-frequency flexor and extensor alternation

Our experimental observations on how perturbations of neuronal activity affect the rhythm of scratching oscillations challenge some of the predictions generated by existing models of CPGs. For example, our finding that ablation of inhibitory V1 or V2b neurons reduces the frequency of scratch oscillations (**Figure 3C, 3D, 3I, S3I**) contradicts the prediction that reducing inhibition should result in faster rhythmic oscillations (*e*.*g*. (Ausborn et al., 2021; Rybak et al., 2006; Sherwood et al., 2011; Skinner et al., 1994)). To address these discrepancies, we developed a novel neuromechanical model capable of dissecting the mechanisms underlying: 1) the decrease in oscillation frequency induced by ablation of excitatory V2a neurons (**Figure 2F, 2G**); 2) the increase in scratching rhythm due to activation of excitatory V2a neurons (**Figure 2I, S2I**); 3) the decrease in oscillation frequency caused by ablation of inhibitory V1 and V2b neurons (**Figure 3C, 3D, 3I, S3I**); and, 4) the atonia driven by activation of V1 neurons (**Figure 3F**). Further, our goal was not only to build a model capable of replicating our experimental results, but also able to generate new predictions on circuit dynamics.

We developed a rate model of the ipsilateral premotor circuit driving flexor and extensor oscillations based on the population-average firing rate and synaptic activity of neuronal populations rather than individual neurons (see **STAR Methods**). We opted for a rate model as it enables simplifying conductance-based models by assuming constant or slowly varying inputs and asynchronous firing (Shriki et al., 2003), thus allowing it to be used to compute circuit dynamics and characterize their underlying mechanisms via simplified schemes (Golomb et al., 2006, 2022; Hayut et al., 2011). The circuit architecture of our rate model (**Figure 4A**) was built on four assumptions. First, the model has two modules, one controlling flexion (F) and one extension (E), coupled via inhibition. Motoneurons in the flexor module (MF) are innervated by excitatory flexor neurons (EF) and inhibitory extensor neurons (IX). Extensor motoneurons (MX) receive mirrored input. Second, the two excitatory neuronal populations (EF and EX) generate rhythmic activity by network bursting. In principle, network bursting can be driven either via intrinsic mechanisms of individual neurons (Coombes and Bressloff, 2005; Husch et al., 2015; Zhong et al., 2010), or by these neurons functioning as a population of coupled excitatory neurons with adaptation currents (Van Vreeswijk and Hansel, 2001). In our model, network bursting is generated via adaptation and intrinsic depolarization currents. Third, there are recurrent intra-module excitatory-inhibitory connections that modulate the frequency of rhythmic bursting. Fourth, excitatory neurons have facilitation mechanisms (Zhong et al., 2010), as facilitation has been shown to modulate the oscillation frequency in networks with bursting neurons (Jackman and Regehr, 2017; Melamed et al., 2008; Reid et al., 1999).

**Figure 4.**
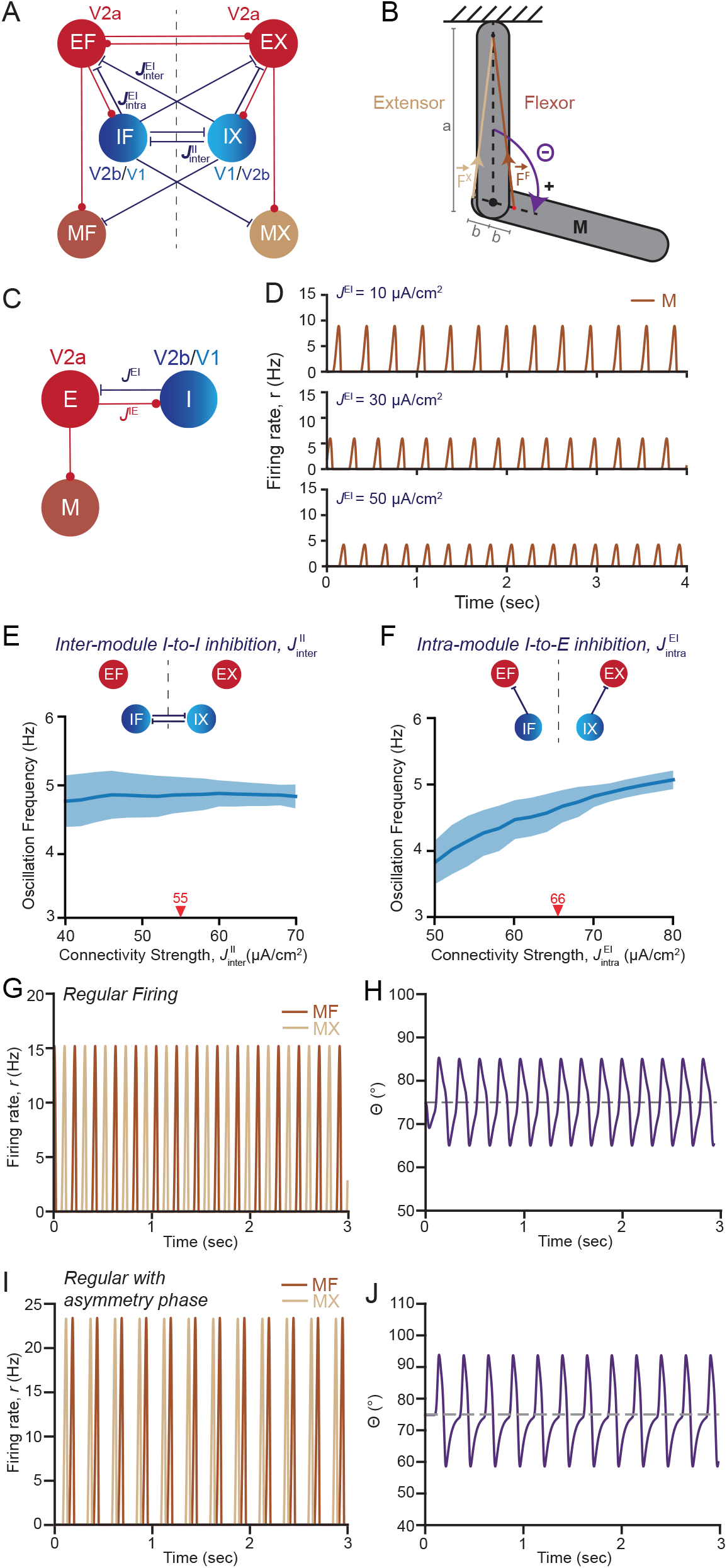
Neuromechanical model for high-frequency flexor and extensor alternation. A) Schematic illustrating the circuit architecture of the neuromechanical model divided into flexor and extensor (X) modules, each comprising excitatory (E), inhibitory (I), and motoneuron populations (M). B) Schematic illustrating the mechanical model of the ankle joint including a static and a rotating segment coupled by flexor and extensor muscles. C) Schematic illustrating an individual module including coupled excitatory (E) and inhibitory (I) neurons and a motoneuron (M) population driven by E. D) Computed dynamics of a single module. Time traces of motoneuron (M) firing rates are plotted for distinct strengths of I-to-E inhibitory conductance *J*^*EI*^. E,F) Simulations of the oscillation frequency generated by the neuromechanical model as a function of the strength of the inter-module I-to-I inhibition 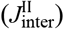 (E) and the intra-module I-to-E inhibition 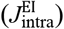 (F). Red arrowheads and numbers indicate the value of synaptic strength used as reference parameter in the rate model (**Table 1**). Data presented as mean ± SD of 50 realizations of synaptic conductance parameters, SD represented as shaded area. G,H) Simulated firing rates of flexor (MF) and extensor motoneurons (MX) (G) and joint angle θ (H) generated by the neuromechanical model using the reference parameter set in **Table 1**. I,J) Simulated firing rates of MF and MX (I) and joint angle θ (J) generated by the neuromechanical model when 20% of V1 neuron-driven inhibition is eliminated (*p*^V1^=0.2).

Our model includes six neuronal populations, two excitatory (EF and EX), two inhibitory (IF and IX), and two motoneuron pools (MF and MX) (**Figure 4A**, see also **STAR Methods**). The EX and EF populations include an equal number of bursting V2a neurons, since V2a neurons do not display a bias in innervating flexor vs extensor motoneurons (Azim et al., 2014; Dougherty and Kiehn, 2010; Hayashi et al., 2018). Since ablation of Chx10-V2a neurons did not completely abolish the scratch rhythm (**Figure 2F, 2G**), the E populations include an additional rhythmogenic excitatory neuron type that we propose is analogous to the Shox2-V2a neurons or the ipsilateral V3 neurons (Dougherty et al., 2013; Zhang et al., 2008). The I populations include V1 and V2b neurons that are not intrinsically bursting but have adaptation currents. The preferential inhibition of flexor motoneurons by V1 neurons and extensor by V2b neurons (Britz et al., 2015) is reflected in the model by the I population innervating flexor motoneurons (IX) being mostly composed of V1 neurons, and the I population innervating extensor motoneurons (IF) preferentially composed of V2b neurons (**Figure 4A, S4A**). Moreover, since V1 neurons are twice as abundant as V2b neurons in the mouse spinal cord (U19-SCC, unpublished data), they account for a higher fraction of I neurons (**Figure S4A**). The strength of synaptic connectivity among the different neuronal populations is represented by the synaptic coupling conductance (*J*), separated into intra-(*J*_intra_) or inter-module (*J*_inter_), and differentiated based on the post- and pre-synaptic neurons (e.g. inhibitory-to-excitatory conductance is denoted as *J*^EI^) (**Figure 4A, STAR Methods**).

### Neuromechanical model of oscillations around a single joint

To relate our model CPG activity with limb movement, we constructed a biomechanical model of the mouse ankle joint. We modelled the ankle joint as a one degree-of-freedom, two-link segment model (**STAR Methods**), in which only the distal segment can rotate around an axis in the fixed proximal segment (**Figure 4B**). The distal segment is actuated by an antagonistic pair (flexor and extensor) of identical muscles, connecting the proximal and distal segments in a bow-string configuration (**Figure 4B**). Next, we used a Hill-type muscle model to convert motoneuron activity into muscle force. This model includes known muscle properties: the low-pass filtering effects of muscle, the length-dependent muscle activation, force-length relationship, and a passive elastic element (**Figure S7, STAR Methods**).

### Increases in inhibition or excitation within a single module drive faster motoneuron bursting

We first analyzed the dynamics of the three neuronal populations (E, I, and M) in a model of a single module (**Figure 4C**). Rhythmic firing of post-synaptic motoneurons (M) is driven by the bursting excitatory population (E) that is activated by an excitatory tonic input (**Figure S4B, S4C**). Firing of inhibitory (I) neurons is also elicited by the E neurons during their active phase and contributes to terminating E neuron bursting (**Figure 4C**). At the end of E neuron bursting, the adaptation current begins to decay, and the latency of its decay is determined by the beginning of the next active phase (**Figure S4D**). Strengthening the I-to-E inhibitory conductance (*J*^*EI*^) decreases the adaptation current, therefore reducing the latency of its decay. This accelerates the termination of excitatory bursting and the beginning of a new active phase, thus effectively increasing the frequency of motoneuron bursting (**Figure 4D, S4E**). Similar increases in frequency were obtained by strengthening the E-to-I excitatory conductance (*J*^*IE*^) (**Figure S4F**). Thus, within a single module, the activity of E and I populations and the strength of their synaptic connections regulate the frequency of motoneuron bursting.

### The differential contribution of intra- and inter-module inhibition to oscillation frequency

Three types of inhibitory synapses were included in our model: 1) inter-module I-to-I inhibition 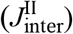 between the IF and IX populations; b) inter-module I-to-E inhibition 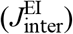 between the IF-EX and IX-EF populations; and, 3) intra-module I-to-E inhibition 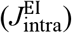 between the IF-EF and IX-EX populations (**Figure 4A**). To examine the contribution of individual inhibitory synaptic conductances to rhythm generation, we determined how the oscillation frequency changes depending on the strengths of individual inhibitory connections. Decreases in inter-module I-to-I inhibition, 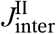, have almost no effect on the oscillation frequency (**Figure 4E**). Reducing the inter-module I-to-E inhibition, 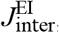, slightly elevates the frequency (**Figure S4G**). Remarkably, reductions in intra-module I-to-E inhibition, 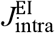, decrease the frequency (**Figure 4F**), consistent with our analyses of the dynamics in a single module model (**Figure 4D**). While reducing the intra-module inhibition increases the rhythm, reductions in inter-module I-to-E inhibition slow it down. Therefore, the effects on frequency caused by reducing inhibition in the entire circuit (ablation efficiency of inhibitory neurons, *p*) will depend on the relative strength of the individual inhibitory synapses. For example, reducing overall inhibition decreases frequency for small 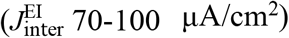 values of inter-module I-to-E inhibition and increases it for larger values 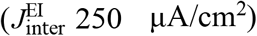 (**Figure S4H)**. Notably, the strength of inter-module I-to-I inhibition does not affect how reductions in overall inhibition impact the oscillation frequency (**Figure S4I**). The analyses of our model suggest that the different inhibitory connections have distinct roles in rhythmogenesis, and the frequency of oscillations is set by the relative strengths of all the individual inhibitory synapses.

In current spinal CPG models based on “half-centers” and “unit bursting generators”, the dominant rhythmogenic inhibitory connections are the ones between the “rhythm generators” (Ausborn et al., 2018; Dougherty and Ha, 2019). Each “rhythm generator” needs at least one slow variable to generate bursting: either an activation variable of a hyperpolarizing conductance or an inactivation variable of a depolarizing conductance (Rinzel and Ermentrout, 1998). Decreasing the inhibitory connections makes each module spend less time in its passive state, causing the slow variable to be larger at the beginning of the next active state, thus, effectively causing an increase in frequency. However, the reduction in frequency caused by the ablation of inhibitory neurons observed in our experiments (**Figure 3C, 3I**) contradicts these predictions. Therefore, we propose that, during scratching, the dominant inhibitory connection is the intra-module I-to-E inhibition, 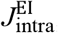, rather than the inter-module I-to-E inhibition, 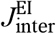, or the inter-module I-to-I inhibition, 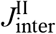. In sum, the strong intra-module I-to-E inhibition in our model is sufficient to explain how decreasing inhibition can effectively reduce the frequency of scratch oscillations (**Figure 4F**).

### Computed motoneuron firing patterns following perturbations of neuron activity

To quantitatively describe our experimental manipulations, we computed the ablation of a population as a decrease in its total synaptic conductance output (see equations 3 and 4 in **STAR Methods**), and the activation as an increase in the input current received (see equations 5 and 6 in **STAR Methods**). The ablation efficiency is defined by the parameter *p* (see equations 3 and 4 in **STAR Methods**), which ranges from 0 (no ablation) to 1 (100% ablation). Activation efficiency is defined by the injected current, *I*_act_, (see equations 5 and 6 in **STAR Methods**). Changing these two parameters to mimic either neuronal ablation or perturbations of their activity varies the dynamical states of the coupled modules, resulting in different patterns of MX and MF activity that range from regular anti-synchronous firing (**Figure 4G**) to constant (100%) co-contraction (**Figure S4L**). In the absence of any ablation (*p=0*), MF and MX fire anti-synchronously (**Figure 4G**), driving oscillations (**Figure 4H**) at similar frequencies to what observed when mice scratch (**Figure 1D**). Low-efficiency ablation of V1 neurons (*p*^V1^=0.2), causes MF and MX to exhibit a phase difference (*ϕ*) between 0 and 2π (**Figure 4I**). This is due to the asymmetrical inhibition exerted by V1 and V2b neurons onto flexor and extensor motoneurons (Britz et al., 2015), which skews MF and MX bursting (**Figure 4I**). Notably, this pattern of regular firing with asymmetric phase retains the ability to generate fluid limb movements (**Figure 4J**). High-efficiency ablation of V1 neurons (*p*^V1^=0.8) causes MF and MX to fire with an irregular, non-periodic pattern, in terms of both frequency and amplitude of oscillations (**Figure S4J, S4K**). High-efficiency ablation of both V1 and V2a neurons (*p*^V1^*=p*^V2a^ =0.95) significantly weakens the inter-module I-to-I and I-to-E inhibition, which causes simultaneous firing of MF and MX, driving a 100% co-contraction state (**Figure S4L**) and restricting the amplitude of joint movements (**Figure S4M**). High-efficiency activation of V1 neurons 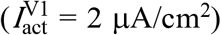 completely suppresses MF and MX activity (**Figure S4N**), causing the limb to settle into an atonia-like state (**Figure S4O**), as experimentally observed following chemogenetic activation of V1 neurons (**Figure 3F**).

### Computed circuit dynamics following manipulations of activity of individual neuronal populations

To mechanistically define the contribution of individual neuronal populations to rhythm generation, we examined how changing parameters in the model to mimic our experimental perturbations modifies the firing pattern, the phase difference, and the oscillation frequency. To mimic in the model the experimental ablation of V2a neurons, we decreased the total synaptic conductance output of the subset of neurons within the E populations that corresponds to V2a neurons (*p*^V2a^). Ablation of V2a neurons in the model (*p*^V2a^) (**Figure 5A**) decreases the oscillation frequency (**Figure 5B, S5A**), in agreement with our experimental results (**Figure 2F,2G**). Since V2a neurons are symmetrically represented in the EF and EX populations, their ablation does not affect the phase difference (*ϕ*) of MF and MX activity, which continues to display a regular firing pattern (**Figure 5B**). Since in our simulations the synaptic conductance output is equally decreased for all E connections, the extent to which ablation of V2a neurons reduces the oscillation frequency depends on the strength of the intra-module I-to-E inhibition 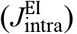. Ablation of V2a neurons in the model causes a reduction in the intra-module E-to-I excitation conductance 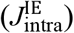 (**Figure S5B**). This decrease in intra-module E-to-I excitation becomes less effective in decreasing the oscillation frequency when the intra-module inhibition is weak 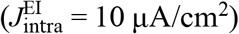. In the absence of intra-module inhibition 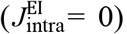, weakening the intra-module E-to-I excitation causes no further reductions in oscillation frequency (**Figure S5B**). Thus, reductions or increases in intra-module excitation require effective intra-module inhibition to affect the oscillation frequency (**Figure 4D, 4F**). The effects of intra-module excitation on the oscillation frequency also depend on the strength of inter-module I-to-E inhibition 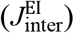. Since reducing inter-module inhibition increases frequency (**Figure S4G**), a very strong 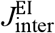 would prevent our model from replicating the experimental observations. Notably, the role of intra-module excitation 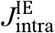 differs from other CPG models, in which the rhythm generating layer includes two excitatory populations that inhibit each other via two intermediate inhibitory populations (Ausborn et al., 2018). In these models, ablation of excitatory neurons, mimicked via reducing the intra-module excitation would result in increases in the bursting frequency.

**Figure 5.**
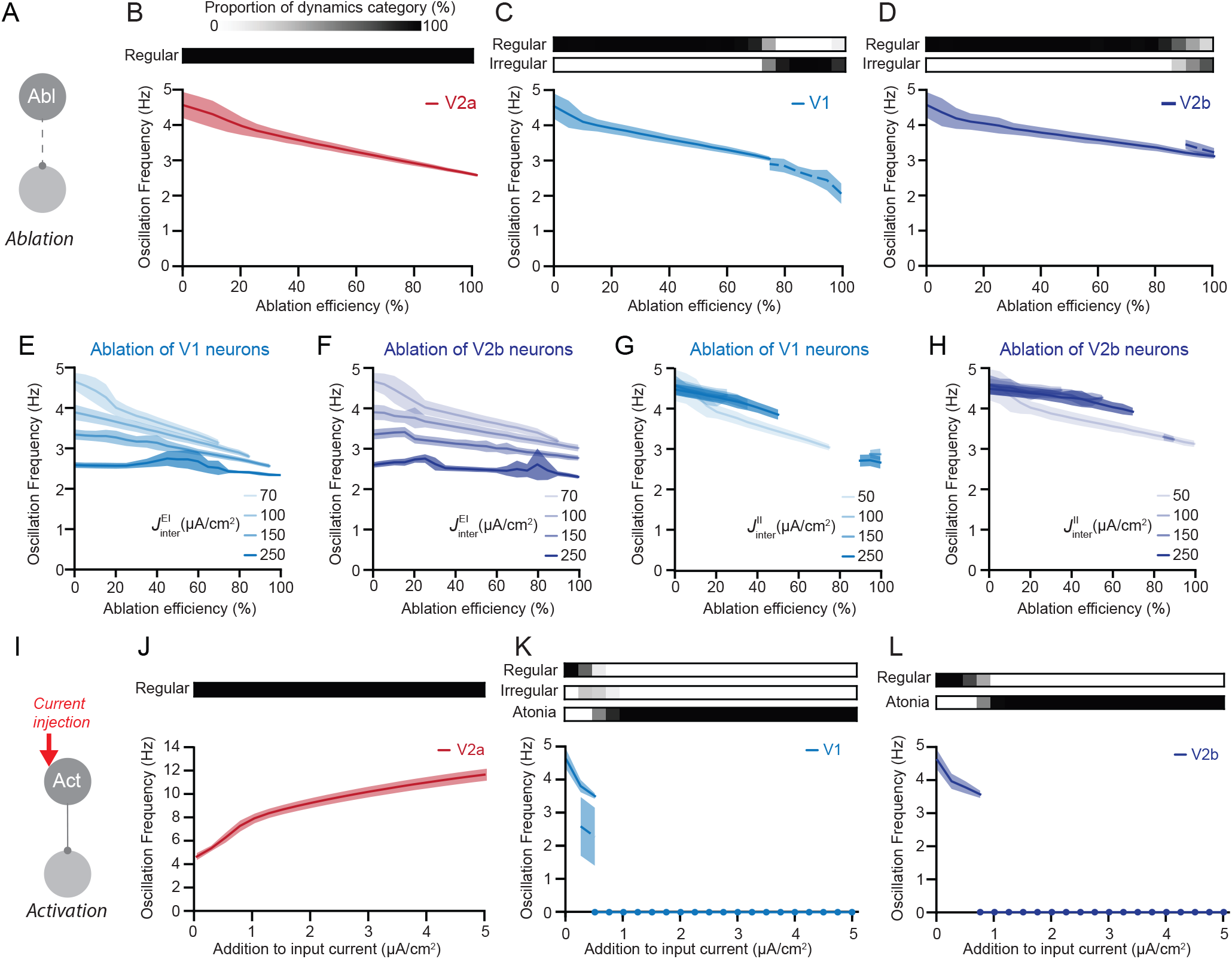
Computed circuit dynamics following manipulations of activity of individual neuronal populations. A,I) Schematics illustrating the simulations of neuronal ablation (A) and activation (I). B-D) Graphs showing the computed oscillation frequency and types of motoneuron firing pattern as a function of the ablation efficiency (*p*) of V2a (B), V1 (C), and V2b (D) neurons. The bars above the graphs represent the occurrence of distinct motoneuron firing patterns (e.g., regular and irregular, see also **STAR Methods**) as a function of the percentage of ablated neurons. E-H) Graphs showing oscillation frequency as a function of the ablation efficiency (*p*) of V1 (E,G) and V2b (F,H) neurons and of the strength of inter-module I-to-E inhibition 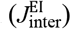 (E,F) and I-to-I inhibition 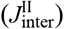 (G,H). Only results for regular oscillations are included in this plot. J-L) Graphs showing the computed oscillation frequency and types of motoneuron firing pattern dependence on the activation current *I*_act_ injected into V2a (J), V1 (K), and V2b (L) neurons. The bars above the graphs represent the occurrence of distinct motoneuron firing patterns as a function of the addition of input current. Solid and dashed lines represent regular and irregular oscillations, respectively. Data presented as mean ±S D of 50 realizations of synaptic conductance parameters, SD represented as shaded area.

Ablation of the subsets of neurons within the I populations corresponding to V1 (*p*^V1^) or V2b (*p*^V2b^) neurons reduces the frequency of oscillations (**Figure 5C, 5D, S5C, S5D**), recapitulating our experimental results (**Figure 3C, 3D, 3I, S3I**). Motoneuron firing is regular at low-efficiency ablation, and becomes progressively irregular (**Figure 5C, 5D**) with larger phase differences as the ablation efficiency of either V1 or V2b neurons increases (**Figure S5E, S5F**). Given the asymmetrical input that these two classes of inhibitory neurons exert on MF and MX, their ablation causes the coupled modules to exhibit large frequency differences that ultimately cause irregular dynamics (**Figure 5C, 5D**) (Schuster and Just, 2005). While V1 neuron ablation has strong effects in causing irregular firing (**Figure 5C**), ablation of V2b neurons has milder ones, with regular and irregular firing states coexisting, even at higher ablation efficiency (**Figure 5D**). The stronger effects induced by ablation of V1 neurons stem from the fact that these neurons are more numerous than V2b neurons and are asymmetrically distributed between the IX and IF populations (**Figure S4A**). Thus, ablation of V1 neurons leads to a stronger reduction in inter-module inhibition and to a higher asymmetry in the number of neurons remaining after ablation in the IX vs the IF population. The strength of inter-module I-to-E inhibition 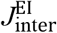 determines the extent to which the ablation efficiency of both V1 and V2b neuron populations affects the oscillation frequency (**Figure 5E, 5F**). The oscillation frequency accelerates as the intra-module inhibition 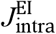 increases (**Figure 4F**) and the inter-module inhibition 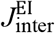 decreases (**Figure S4G**). Since ablation of V1 or V2b neurons reduces inhibition in the entire circuits, decreasing both intra- and inter-module inhibition, the net effect on frequency depends on the relative strength of individual inhibitory synapses. Ablation of V1 or V2b neurons causes small reductions in oscillation frequency for strong inter-module I-to-E inhibitory conductance 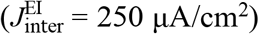, and larger decreases for weaker values 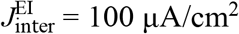 (**Figure 5E, 5F**). Since the inter-module I- to-I inhibition 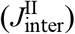 does not affect the oscillation frequency (**Figure 4E**), changes in its strength do not modulate how V1 or V2b neuron ablation efficiency impacts frequency (**Figure 5G, 5H**).

Activation of the V2a subset within the E populations (**Figure 5I**) increases the oscillation frequency (**Figure 5J, S5G**), as also shown in our experimental results (**Figure 2I, S2I**). This increase is due to facilitation mechanisms in the E neurons (**Figure S5H-S5J**). Low-efficiency activation of V1 neurons 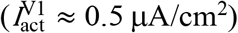 causes irregular MF and MX activity, as the inhibition exerted by the two modules becomes asymmetric (**Figure 5K**). High-efficiency activation of V1 or V2b neurons 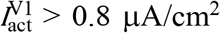 and 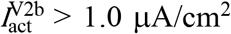 effectively silences the MF and MX populations, causing the circuit to reach a fixed point that represents an atonia-like state (**Figure 5K, 5L, S5K, S5L**), again confirming our experimental results.

In summary, our neuromechanical model not only recapitulates the results of our experimental manipulations but also predicts how perturbations of individual populations affect motoneuron firing pattern and phase differences (**Figure 5B-5D, 5J-5L**). Moreover, the model reveals the relative contribution that individual synaptic connections make to set the rhythm of oscillation (**Figure 5E-5H, S5B**) and predicts the threshold at which activation of V1 and V2b inhibitory neurons drives atonia (**5K, 5L, S5K, S5L**).

### Distinct cooperation dynamics among the ipsilateral neuron populations in driving oscillations frequency

Finally, we experimentally investigated how excitatory and inhibitory populations cooperate to drive the rhythm of scratch oscillations using our genetic approach to simultaneously manipulate the activity of excitatory V2a and inhibitory V2b neurons (*Hes2*^*iCre*^;*hCdx2::FlpO*) (Hayashi et al., 2023) or inhibitory V1 and excitatory V2a neurons (*En1*^*Cre*^;*Chx10*^*Cre*^;*hCdx2::FlpO*).

The simultaneous ablation of excitatory V2a and inhibitory V2b neurons (**Figure 6A, S6A**) reduced the frequency of scratch oscillations (**Figure 6B**) and the number of scratch bouts (**Figure S6B**) to the same extent as ablation of only V2a neurons. We then simulated this dual manipulation in our model, recapitulating the reduced frequency (**Figure 6C, S6C, S6D**). When V2a and V2b neurons are ablated with high efficiency (*p*^V2a^ = *p*^V2b^ = 0.95), MX and MF fire in irregular patterns (**Figure 6C**), as observed following the ablation of V2b (**Figure 5D**) but not V2a neurons (**Figure 5B**). Intriguingly, when the V2a and V2b neurons are simultaneously removed, at an intermediate level of ablation efficiency (*p*^V2a^ = *p*^V2b^ = 0.7), an irregular pattern, which is not seen when V2a or V2b interneurons are individually ablated, emerges (**Figure 6C**). In sum, V2a and V2b neurons have partly redundant roles in the circuit, as the decrease in oscillation frequency induced by their simultaneous ablation is not stronger than the frequency reduction caused by the individual ablation of V2a neurons (**Figure 6B**).

**Figure 6.**
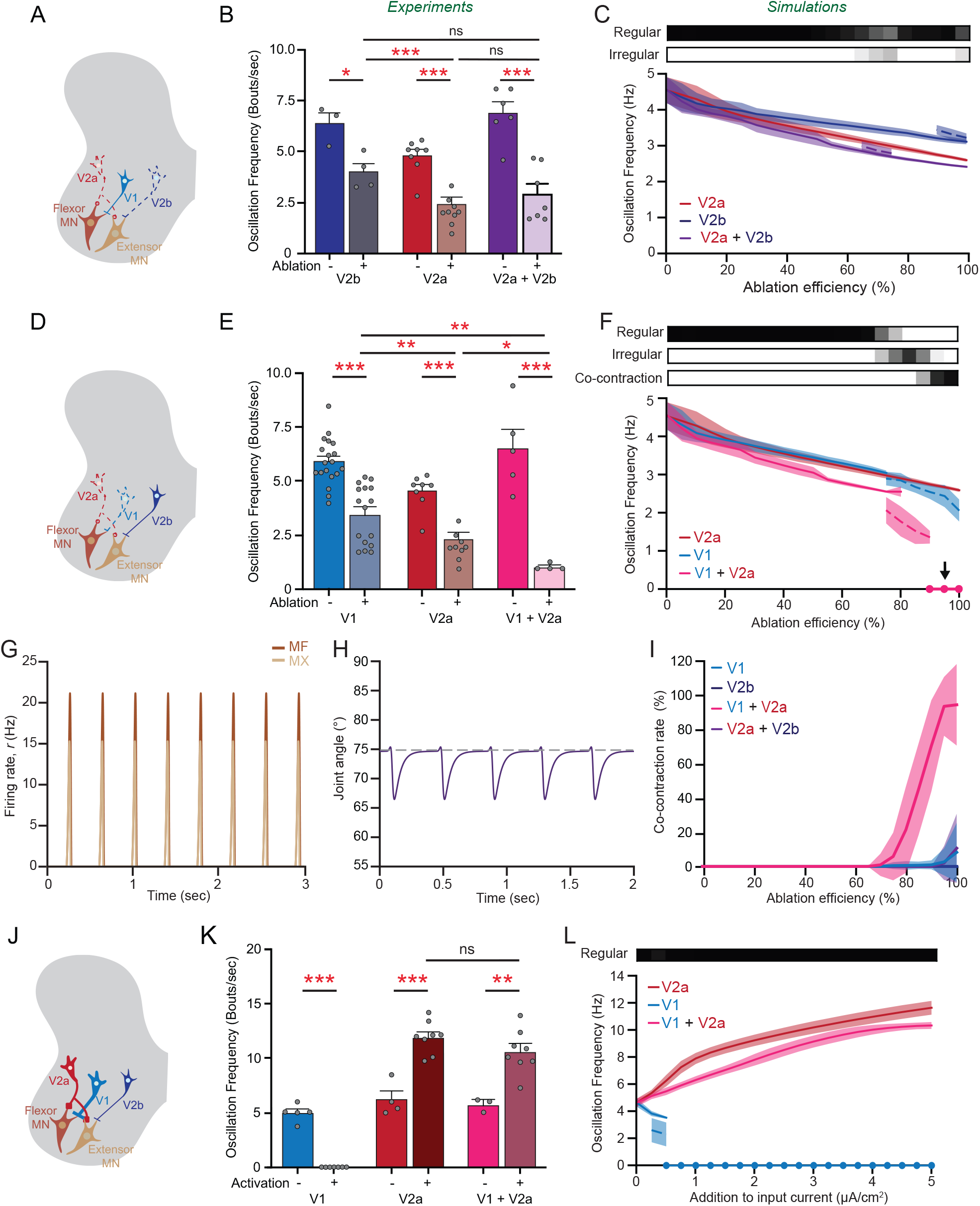
Distinct cooperation dynamics among the ipsilateral neuron populations in driving oscillations frequency. A,D,J) Schematics illustrating the ablation of V2a and V2b neurons (A), the ablation of V1 and V2a neurons (D), and the CNO-driven activation of V1 and V2a neurons (J). B) Bar graph showing the reduction in scratching oscillation frequency in V2a neuron-ablated, V2b neuron-ablated, and dual V2a and V2b neuron-ablated mice compared to littermate controls, p = *0*.*0002* V2a and V2b control vs ablated, *p* = *0*.*0009* V2b vs V2a ablated, *p* = *0*.*1719* V2b vs V2a and V2b ablated, *p* = *0*.*1270* V2a ablated vs V2a and V2b ablated. C) Graphs showing the computed oscillation frequency and types of motoneuron firing pattern as a function of the ablation efficiency (*p*) of V2b, V2a, and both V2a and V2b neurons. E) Bar graph showing the reduction in scratching oscillation frequency in V1 neuron-ablated, V2a neuron-ablated, and dual V1 and V2a neuron-ablated mice compared to littermate controls, *p* = *0*.*0003* V1 and V2a control vs ablated, *p* = *0*.*0057* V1 vs V2a ablated, *p* = *0*.*0027* V1 vs V1 and V2a ablated, *p* = *0*.*0151* V2a ablated vs V1 and V2a ablated. F) Graphs showing the computed oscillation frequency and types of motoneuron firing pattern as a function of the ablation efficiency (*p*) of V1, V2a, and both V1 and V2a neurons. The arrow points to the value *p* = 0.95 (panels G,H). G,H) Simulated time traces of motoneuron firing rate (G) and joint angle (H) when V1 and V2a neurons are ablated with high-efficiency (above 90%), causing a constant co-contraction state. This state is not considered as an “oscillation” in panel F because the movement does not exceed the threshold (5°; see **STAR Methods**) in the extensor direction. I) Graphs showing that the 100% co-contraction state occurs following high-efficiency ablation of V1 and V2a neurons but not following other manipulations. K) Bar graph showing the atonia in V1 neuron-activated mice, and the increase in frequency following V2a and V1 and V2a neuron activation, *p < 0*.*0001* V1 control vs activated, *p* = *0*.*0028* V1 and V2a control vs activated, *p* = *0*.*1596* V2a activated vs V1 and V2a activated. L) Graphs showing the computed oscillation frequency and types of motoneuron firing pattern as a function of the injected current (*I*) in V1, V2a, and both V1 and V2a neurons. Experimental data presented as mean ± SEM, individual mice represented as filled grey circles, SEM as shaded area. Statistical analysis performed using two-tailed Student’s *t-test*. Modeling data presented as mean ± SD of 50 realizations of synaptic conductance parameters, SD represented as shaded area. The bars above the line graphs in panels C, F, and L represent the occurrence of distinct motoneuron firing patterns (e.g. regular, irregular) as a function of the ablation efficiency (C,F) or the addition to input current (L).

Simultaneous ablation of excitatory V2a and inhibitory V1 neurons (**Figure 6D, S6E**) strongly reduced the ability of mice to scratch (**Figure S6F**), with oscillation frequency falling to 1 Hz (**Figure 6E**). Simulating this perturbation in our model reproduces the experimental data (**Figure 6F, S6G, S6H**), and shows a 100% co-contraction state at high-efficiency ablation (**Figure 6F, 6G, 6I**). This 100% co-contraction state is observed when V2a neurons are co-ablated with V1 neurons, but not V2b neurons (**Figure 6I**), likely because V1 neurons are more abundant than V2b neurons (**Figure S4A**). Together, these results suggest that V1 and V2a neurons function in synergy to control the oscillation frequency.

Finally, we asked if the co-activation of excitatory V2a and inhibitory V1 neurons results in faster oscillations, as seen with V2a neuron activation (**Figure 2I, S2I**), in atonia, as caused by V1 neuron activation (**Figure 3I**), or in a different response due to distinct cooperation dynamics. The simultaneous activation of V1 and V2a neurons (**Figure 6J, S6I**) caused an increase in the frequency of scratch oscillations (**Figure 6K**) to almost the same extent (slightly smaller but not significantly different) as the activation of V2a neurons alone. No changes in the total number of scratch bouts were observed (**Figure S6J**). Simulating this perturbation in our model shows similar results, causing an increase in oscillation frequency that is somewhat smaller than the effect of activating only V2a neurons (**Figure 6L, S6K, S6L**). The simultaneous activation of both V1 and V2a neurons does not induce irregular firing nor atonia (**Figure S6L**). Our model also predicts that simultaneous activation of excitatory V2a and inhibitory V2b neurons (**Figure S6M**) will similarly affect the frequency of oscillations and not be stronger than what is observed by activating V2a neurons only (**Figure S6N, S6O**).

In summary, we provide experimental and computational evidence that distinct circuit dynamics underlie the cooperation among the excitatory V2a and the inhibitory V1 and V2b neuron populations in controlling rhythm and phase of motoneuron firing.

## Discussion

We have experimentally defined the key contributions of three cardinal classes of spinal ipsilaterally projecting neurons, the excitatory V2a and the inhibitory V1 and V2b neurons, to scratch rhythm generation. The reduction in scratch oscillation frequency caused by the loss of inhibitory V1 and V2b neurons (**Figure 3C, 3D, 3I, S3I**) contradicted predictions from other CPG models, like the classical “half-center” (Brown, 1911, 1914; Skinner et al., 1994) and the “unit burst generator” (Ausborn et al., 2018; Dougherty and Ha, 2019), in which decreasing inhibition results in increases in network bursting frequency. To address this discrepancy and mechanistically explain our experimental data, we built and analyzed a neuromechanical model, in which inhibitory-coupled flexor and extensor modules drive oscillations of a single joint. Using experimental and modeling approaches, we showed that non-linear cooperation dynamics underlie the contribution of excitatory and inhibitory populations to rhythm and phase of motoneuron firing. Our biomechanical model not only recapitulates our experimental observations but also makes predictions about the role of each synaptic conductance and circuit dynamics in the physiological and perturbed execution of the scratch reflex.

### Model hypotheses and their effects on neural dynamics

Analyses of our model in physiological and perturbed conditions showed that three conditions are needed for scratch-like oscillations to emerge and be altered in a way that mimics the responses to experimental manipulations of neuronal activity: a) each module should oscillate independently (**Figure 4C), b**) intra-module inhibition should be stronger than a critical threshold, whose value increases depending on the strength of the inter-module inhibition (**Figure S4H**), and c) synapses from excitatory neurons should have facilitation mechanisms (**Figure S5H**).

First, we propose that the E populations act as self-bursting networks. Individual modules oscillate independently via an intrinsic mechanism in E neurons that involves two ionic currents, one for depolarization and one, slower, for adaptation (**Figure S4D, S4E**). This intrinsic mechanism is of course not the only possibility for generating self-bursting networks. Alternative mechanisms include network bursting due to tonically firing excitatory neurons with adaptation currents (Van Vreeswijk and Hansel, 2001) or to excitatory neurons recruiting inhibitory neurons via facilitation mechanisms (Hayut et al., 2011; Melamed et al., 2008). Future electrophysiological recording experiments discriminating between these scenarios and defining the contribution of the other excitatory populations will help to clarify the mechanisms underlying network bursting.

Second, the discrepancies on the role of inhibition in modulating rhythm between the predictions of other CPG models (Ausborn et al., 2018; Golomb et al., 2022; Soloduchin and Shamir, 2018) and our experimental data prompted us to evaluate the contribution of individual inhibitory synapses to rhythmogenesis. Decreasing the intra-module inhibition reduces the bursting frequency (**Figure 4F**), whereas decreasing the inter-module I-to-E inhibition increases frequency (**Figure S4G**). Therefore, to replicate our experimental results, reductions in the overall inhibition should weaken more intra-module than inter-module inhibition (**Figure S4H**). The predicted differences in synaptic strength of the distinct inhibitory synapses need to be validated using *in vitro* and *in vivo* recordings. An alternative hypothesis could be that there are specialized subsets within the V1 and V2b populations that differentially contribute to intra-module and inter-module inhibition. Targeted anatomical tracings and manipulations of specific V1 neurons subsets (Bikoff et al., 2016; Trevisan et al., 2024), will start to reveal whether the different roles of the inhibitory connections between the I and E populations predicted by the model arise from different synaptic connectivity strenghts, distinct subsets of V1 and V2b neurons, or both mechanisms.

Third, to explain the increase in scratching frequency observed following activation of V2a neurons (**Figure 2I, S2I**), our model hypothesizes the presence of facilitation mechanisms in E neuron synapses. Activation of V2a neurons increases the firing rate and, via facilitation (**Figure S5H**), increases the effective intra-module excitation that accelerates scratch oscillations (**Figure S4F**). An alternative solution to facilitation mechanisms is that the activation-driven increase in bursting is due to intrinsic neuronal properties, as shown in other models (e.g. (Golomb et al., 2006)), or is due to a combination of intrinsic properties and E-to-E synaptic connections (Van Vreeswijk and Hansel, 2001). In adult mice, serotonin decreases membrane input resistance in V2a neurons, facilitating evoked plateau potentials (Husch et al., 2015). Whether the increase in oscillation frequency is due to facilitating synapses, intrinsic mechanisms in V2a neurons, or a combination of synaptic and intrinsic neuronal properties remains to be assessed.

### Comparison with cell type-agnostic models for rhythmic scratching

We developed a novel rate model, in which rhythm and phase of antagonist motoneuron firing are tightly linked in a single layer. In this model the rhythm is driven by a self-bursting population of excitatory neurons (E), while inhibitory neurons (I) have a dual function, i.e. controlling bursting frequency via the intra-module and inter-module I-to-E inhibition (**Figure 4F, S4G**) and the phase of flexor and extensor motoneuron firing via inter-module I-to-E and I-to-I inhibitory connections (**Figure 4G-4J**). Our approach addresses important gaps in the computational models used so far to describe scratching by providing a mechanistic explanation of how key CPG neuron populations contribute to determining the scratch rhythm and assessing in detail how the relative strength of intra- and inter-module inhibitory conductance affects the oscillation frequency. The robustness of our model with respect to recapitulating the effects of experimental perturbations was also tested by manipulating the activity of individual CPG neuron populations or combinations thereof.

A balanced E-I neural network has been proposed for the generation of the scratch rhythm in turtles (Berg et al., 2007), with neural activity resembling “rotational dynamics” at the population level (Lindén et al., 2022). In the rotational dynamics model, synaptic coupling (Lindén and Berg, 2021; Lindén et al., 2022) is strong (as defined in (Pehlevan and Sompolinsky, 2014; Van Vreeswijk and Sompolinsky, 1996)), random, and sparse, with the model exhibiting the hallmarks of a fluctuation-driven regime (Berg, 2017). Population frequency is modulated by adjusting the gain of selected “speed” or “break” neuronal subsets that are a mix of excitatory and inhibitory neurons. However, in contrast to our model, the rotational dynamics model does not differentiate between individual interneuron populations or predict how they contribute to rhythm and pattern generation. Moreover, its robustness in recapitulating experimental perturbations remains to be assessed. An intriguing future challenge would be to convert our rate model into a spiking, strongly coupled neural model, similar to the rotational dynamic model, while preserving our network architecture and the strong intra-module inhibitory conductance. This will enable us to assess the irregular temporal dynamics of interneuron input onto motoneurons, and how this input determines motoneurons firing frequence and phase in physiological state and following perturbations of neuronal activity.

A more recent version of the rotational dynamics model includes two sparse, randomly connected networks coupled via mutual inhibition, describing the behavior of individual neurons within a rhythm-generating population (Strohmer et al., 2024). In this model, excitatory neurons possess a slow current that enables bursting only in a subset of neurons (Strohmer et al., 2024). The intra-module inhibition widens the range of firing phases of each neuronal population (Strohmer et al., 2024), making the distribution of firing phases closer, but not equal, to what is predicted by the rotational dynamics model (Lindén et al., 2022). Increases in intra-module inhibition cause firing irregularities (a main hallmark of fluctuation-driven regimes), as observed in models of two coupled inhibitory populations (Golomb et al., 2022). However, in this model the relative contribution of intra- and inter-module inhibition to rhythmogenesis was not assessed.

### Cooperation of excitatory and inhibitory neurons in driving rhythm of scratch oscillations

Our results reveal a number of interesting interactions between the V1, V2a and V2b populations with respect to rhythm and pattern generation. While the ablation of V1 and V2b neurons individually results in hyper-flexion and hyper-extension, respectively, which matches their anatomical asymmetry in innervation of flexor and extensor motoneurons (Britz et al., 2015), repeating these ablation experiments together with V2a neuron removal painted a more complex picture. Co-ablating V1 and V2a neurons severely impairs scratch movements (**Figure 6E, S6E**), in line with our model predicting the loss of both causes a 100% co-contraction state (**Figure 6F-6I**). This was not observed following the co-ablation of V2a and V2b neurons (**Figure 6B, 6C, 6I**). Similarly, ablation of V2a neurons rescues the leg hyper-extension phenotype observed in V2b neuron-ablated mice (Britz et al., 2015; Hayashi et al., 2023), whereas it severely worsens the coordination defects in flexor and extensor alternation observed in V1 neuron-ablated mice during locomotion (Gatto, Goulding unpublished data). Our behavioral and computational results show that V2b neurons have a complementary role to V1 neurons with regards to flexor and extensor specificity, but also suggest redundant functions at the circuit level, with the remaining V1 neurons able to compensate for the loss of V2b neurons (**Figure 6B**). This functional redundancy might simply be due to the higher number of V1 neurons in the cord (**Figure S4A**), which we included in our model. Alternatively, it might be due to differences in the intrinsic properties and/or connectivity of these two populations. We know for example that V1 and V2b neurons receive distinct local, descending, and sensory inputs.

Another important aspect that is not addressed in our rate model is the interneuron spike timing and the timing with which excitatory and inhibitory inputs reach motoneurons. In turtles, a balanced increase in excitatory and inhibitory neuron activity has been modeled to drive faster scratching movements (Berg et al., 2007). Similarly, in larval zebrafish, faster swimming is associated with the increased coincidence of excitatory and inhibitory input onto motoneurons (Kishore et al., 2014). Whether the timing of V1 and V2b neuron inputs onto motoneurons in mice follows distinct patterns in relation to V2a neuron input represents an intriguing hypothesis but remains to be assessed. Recording interneuron activity in scratching animals alongside the development of spiking models can help to address these important questions.

### Divergence or convergence of spinal circuits for fast movements

CPGs have been shown to generate diverse motor behaviors by modulating the timing and phase of motoneuron firing (Büschges et al., 2008; Friedman et al., 2009; Marder and Bucher, 2001; Marder and Calabrese, 1996). Microcircuits within the CPG networks are thought to be responsible for the emergence of diverse rhythms, with distinct rhythmic behaviors in turtle and cat recruiting specialized subsets of spinal interneurons (Berkowitz, 2010; Berkowitz and Hao, 2011; Frigon and Gossard, 2010), and swimming speeds in adult zebrafish driven by distinct subsets of interneurons and motoneurons (Ampatzis et al., 2014; Ausborn et al., 2012; Kimura and Higashijima, 2019; Song et al., 2018, 2020). To which extent this cellular and circuit logic is conserved in mice remains to be assessed.

The activation of excitatory Chx10-V2a neurons accelerates scratching movements in mice (**Figure 2I, S2I**). While Chx10-V2a neurons are rhythmically active in fictive locomotor preparations (Dougherty and Kiehn, 2010; Zhong et al., 2010), their removal does not impact locomotor rhythm in behaving mice (Crone et al., 2009). By contrast, the Shox2-V2a neurons are necessary to set locomotor rhythm (Dougherty et al., 2013). As fictive scratch is ∼2.3 times faster than fictive locomotion (Frigon and Gossard, 2010), an intriguing hypothesis is that scratch and locomotion in mouse, similar to swimming at different speeds in zebrafish, require two distinct subsets of V2a neurons – the Shox2-V2a neurons for slow locomotor rhythm and the Chx10-V2a for faster oscillations.

The V1 neurons also contribute to fast rhythmic activity during both locomotion and scratching. Ablating the V1 neurons in zebrafish impairs fast swimming due to aberrant recruitment of slow motoneurons (Kimura and Higashijima, 2019). In mouse, V1 neurons contribute to fast rhythm generation *in vitro* (Gosgnach et al., 2006), and we have also shown that they contribute to the scratch oscillation frequency (**Figure 3B-3D**). Taken together, these findings imply that inhibitory V1 and excitatory V2a might form a conserved circuit motif that is used to drive rhythmic movements, with distinct subsets of V2a neurons contributing to slow vs fast rhythms and inhibitory V1 supporting faster oscillations.

On the other hand, V2b neurons are a conundrum. In zebrafish, suppressingV2b neuron activity increases the tail beat frequency (Callahan et al., 2019), whereas in mice ablating V2b neurons reduces the oscillation frequency (**Figure 3H, 3I**). This raises the question as to whether V2b neurons have acquired a novel function or become a more specialized subset in limbed animals? Future studies comparing V2b neuron heterogeneity and connectivity between zebrafish and mouse circuits should shed light on these hypotheses and might reveal novel configurations of the network architecture.

In conclusion, our findings raise the intriguing theory that the speed-dependent microcircuits identified in zebrafish might also exist in rodents for selected behaviors, suggesting an evolutionary conserved strategy by which fast oscillations of a single joint might be driven by similar dynamics as those seen for undulatory movements in aquatic animals.

## Acknowledgement

We thank members of the NIH U19 - Spinal Circuits for the Control of Dexterous Movement for helpful discussions on experiments and analyses. This work was funded by the National Institutes of Health Brain Initiative (NIH-U19NS112959 to MG, DG, GG, TS and EA, NIH-R01NS111643 to MG, NIH-RF1NS128898 to EA, NINDS-F32NS126231 to AN), the German Research Foundation (CRC1451, Project ID 431549029-A09, Z02 and iBehave Network Grant to GG), and the Israel Science Foundation (grant No. 1511/24 to DG).

## Author contributions

<u>Conceptualization</u>: EA, MG, TS, DG and GG; <u>Methodology</u>: MY, AN, TS, DG and GG; <u>Investigation</u>: MY, AN, DG and GG; <u>Writing Original Draft</u>: MY, DG and GG; <u>Review & Editing</u>: MY, AN, EA, MG, TS, DG and GG; Funding Acquisition, EA, MG, TS, DG and GG; Supervision: EA, MG, TS, DG and GG.

## Declaration of interest

The authors declare no competing interests

## Star ⋆ methods

### Key resources table

| REAGENT or RESOURCE | SOURCE | IDENTIFIER |
| --- | --- | --- |
| Chemicals, Peptides, and Recombinant Proteins |  |  |
| Chloroquine | Sigma | C6628 |
| Clozapine N-oxide (CNO) | Sigma | C0832 |
| Diphtheria Toxin (DTx) | List Biological Lab | #150 |
| Saline Solution | APP Pharmaceuticals | NDC63323-186-10 |
| Experimental Models: Organisms/Strains |  |  |
| Mouse: *Chx10*<sup>Cre</sup> | (Azim et al., 2014) |  |
| Mouse: *hCdx2::FlpO* | (Bourane et al., 2015a) |  |
| Mouse: *En1*<sup>Cre</sup> | (Sapir et al., 2004) |  |
| Mouse: *Hes2*<sup>iCre</sup> | (Hayashi et al., 2023) |  |
| Mouse: *R26*<sup>ds-hM3Dq</sup> | Jackson Laboratory | #026942 |
| Mouse: *R26*<sup>ds-hM4Di</sup> | (Bourane et al., 2015b) |  |
| Mouse: *Tau*<sup>ds-DTR</sup> | (Duan et al., 2014) |  |
| Software and Algorithms |  |  |
| Adobe Creative Cloud | Adobe | www.adobe.com |
| Excel-Office 2018 | Microsoft | www.microsoft.com |
| Image J |  | imagej.nih.gov/ij |
| Prism | GraphPad | www.graphpad.com |
| R | The R project | r-project.org |
| Python |  | https://www.python.org |
| SIMI Motion | SIMI | http://www.simi.com/en/ |

## Resource availability

### Lead Contact

Further information and requests for resources and reagents should be directed to and will be fulfilled by the Lead Contact, Graziana Gatto.

### Materials Availability

All published reagents and mouse lines will be shared upon request within the limits of the respective material transfer agreements.

### Data and Code Availability

All original code will be deposited and publicly available as of the date of publication on GitHub. Any additional information required to re-analyze the data reported in this paper is available from the lead contact upon request.

### Experimental model and subject details

Mice were maintained following the protocols for animal experiments approved by the IACUC of the Salk Institute for Biological Studies according to NIH guidelines for animal experimentation, and by the local health authority in North Rhein Westphalia (LANUV). 6- to 10-week-old mice of both sexes were used for behavioral experiments. Analysis of the behavioral data showed similar responses in male and female mice.

## Methods

### Neuronal Ablation

Mice carrying the *En1*^*Cre*^;*hCdx2::FlpO, Chx10*^*Cre*^;*hCdx2::FlpO, Hes2*^*iCre*^;*hCdx2::FlpO, En1*^*Cre*^;*Chx10*^*Cre*^;*hCdx2::FlpO* alleles in addition to the effector allele *Tau*^*ds-DTR*^ received i.p. injections of diphtheria toxin (DTx, 50 ng/gram of weight; List Biological Laboratories) at P28 and P31. *Hes2*^*iCre*^;*VGAT*^*FlpO*^;*Tau*^*ds-DTR*^ mice and littermate controls *Hes2*^*iCre*^;*Tau*^*ds-DTR*^ mice received intrathecal injections of diphtheria toxin (DTx, 10 ng; List Biological Laboratories) at P28, P30 and P32. *FlpO* negative mice injected with DT were used as controls.

### Drug Administration

Chloroquine was dissolved in 0.9% sterile saline and injected subcutaneously in the nape at a final concentration of 200 *µ*g. Clozapine-N-oxide (Sigma) was dissolved in DMSO and then diluted with 0.9% sterile saline such that the concentration of DMSO did not exceed 1% of the injected solutions. For chemogenetic silencing (*R26*^*ds-hM4Di*^) and activation (*R26*^*ds-hM3Dq*^), experimental mice received i.p. injections of CNO (2 mg/kg of weight). *FlpO* negative mice injected with CNO were used as controls.

### Chloroquine-induced scratching

Mice were acclimatized to the plexiglass chamber for 1 hour for three consecutive days. On the experimental day, 200 *µ*g of chloroquine was injected subcutaneously in the nape using a 0.5 ml insulin syringe with a 29G1/2 needle (Exel). Animals were placed in 6 cm x 6 cm x 8 cm Plexiglas boxes and video was recorded (Panasonic SDR-S26) for 30 minutes after injection at 30 fps. A single scratch bout was defined as every time the mouse hindlimb executed a full circular trajectory around the nape. Videos were blindly scored by the experimenter, and for each scratch episode, the time of occurrence, the duration, and number of bouts were recorded. Raster plots were generated in R using the *ggplot* package. For kinematic reconstructions, mice were injected with 200 *µ*g of chloroquine and placed in 6 cm x 6 cm x 8 cm Plexiglas boxes for video recording using a high-speed camera (mV Blue Cougar XD; 200 frame/second). We collected 5-6 videos of 5-11 sec for each mouse, sampling only a small part of the behavior, and tracked hindlimb 2D kinematics using SIMI Motion only in the videos when the scratching movement occurred in front of the camera and all the hindlimb joints/angles were visible.

### Neuronal activity manipulations and behavioral testing

6- to 10-week-old mice of both sexes were used for behavioral testing following ablation or chemogenetic manipulations of neuronal activity. For ablation experiments, mice were tested 2 weeks after the first DTx injection. For CNO-induced silencing and activation, 6- to 7-week-old mice were tested 10 and 20 minutes following i.p. injection of the drug, respectively. All tests were conducted in the morning to minimize circadian-induced variability in response. The experimenter was blind to the genotype of the animals during the recording and the analysis of the videos and manually annotated the beginning and end of each scratch episode and the number of oscillations (bouts) of each episode. These data were used to build frequency distribution plots of the oscillations for each mouse, which were then averaged for each genotype. The kinematic trajectories were reconstructed using the pattern recognition tracking option in SIMI Motion while manually correcting the mis-tracked points, and joint coordinates were used to derive the ankle angle changes.

### Statistical Analysis

All statistical analyses were done in Prism. Depending on the number of groups and variables to compare, we used one-way ANOVA with Dunnett’s post-hoc test, two-way ANOVA with Bonferroni’s post hoc test, or two-tailed Student’s t-test (paired or unpaired according to the experimental group) as indicated in the figure legend.

## Biophysical Neuronal Rate Model

### Neuronal populations and synaptic conductance strengths

We build a rate model [Shriki et al. (2003), Hayut et al. (2011), Golomb et al. (2022)] of a spinal cord circuit composed of interneurons and motorneurons. We assume that the neuronal populations are segregated into flexor (F) and extensor (X) modules. Each module includes an excitatory population (E), composed mostly of V2a neurons; an inhibitory population (I), composed of V1 and V2b neurons, and a motoneuron population (M). Each population is denoted by its neuronal type (first letter) and its module (second letter), and there are 6 populations: EF, IF, MF, EX, IX and MX (Figure 4A). Commissural interneuron populations are not included in the model as scratch is executed as an unilateral hindlimb movement. Anatomical and physiological evidence show that V1, V2a and V2b interneurons and motoneurons are synaptically connected [Sengupta and Bagnall (2023)], although the strengths of the synaptic connections among the populations remains unknown. There is limited information on the connectivity between excitatory interneurons in one module and antagonist motoneurons, but it is well established that flexor and extensor inhibitory neurons synapse onto extensor and flexor motoneurons, respectively [Britz et al. (2015)]. We neglect proprioceptive input because deafferentation (removing the sensory feedback) did not affect the scratching frequency [Deliagina et al. (1975)]. The conductance *J*^*iα*;*jβ*^ denotes the synaptic coupling strength from the pre-synaptic population *jβ* to the post-synaptic population *iα*, where *α, β ∈* {F, X} and *i, j ∈* {E, I, M}. For example, *J* ^EX;IF^ is the synaptic conductance strength from the IF population to the EX population. Based on published results [Dougherty and Kiehn (2010)], we assume the excitatory conductance strength from excitatory populations (EF and EX) to be symmetrical between flexors and extensors. Under this assumption, the index for flexor/extensor can be omitted. We denote whether the conductance is “inter-” or “intra-” the flexor/extensor modules. The conductance from population EF and EX is denoted for *α, β ∈* F, X by

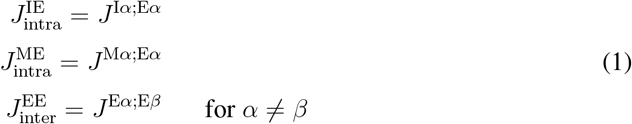

Similarly, we assume flexor-extensor symmetry between the inhibitory conductance strengths and define:

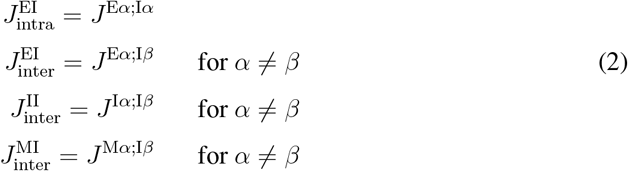

We consider that the excitatory neuronal populations is composed mostly by V2a neurons (proportion being *κ*^E^ = 0.8), and also includes a small fraction of either neurons, analogs of Shox2+ neurons [Dougherty et al. (2013)] and/or V3 neurons [Zhang et al. (2008)]. For inhibitory populations, we consider that V1 and V2b neurons differ in absolute numbers and innervate flexor and extensor motoneurons asymmetrically [Britz et al. (2015)]. Specifically, V1 neurons are twice as many as V2b neurons (Silverman, Goulding unpublished data) and the fraction of V1 neurons is larger than the fraction of V2b neurons among the interneurons that inhibit flexor motoneurons, and vice versa for extensor (Figure S4A). This asymmetry in anatomy resulted in asymmetrical impairments of the duty cycle of flexor and extensor muscles following ablation of V1 and V2b neurons [Britz et al. (2015)]. We model V1 and V2b neurons as a single I population for both flexor and extensor modules. This simplification aims to reduce the complexity of this nonlinear system, as well as to focus on a minimal circuit that could recapitulate the experimental results. In this simplified network, we model the asymmetry of V1/V2b by introducing scaling parameters. We define parameters *κ*^I*α*^ to represent the relative number of V1 neurons in an *α*-th inhibitory population, with the relative value of V2b neurons being 1-*κ*^I*α*^.

### Modification of synaptic conductance strength during neural ablation and activation

Neural ablation is modeled as a reduction in the strength of the synaptic connections from a specific population. We denote the ablation efficiency of a population by *p* with the superscript indicating the population: V2a, V1 or V2b; *p ∈* [0, 1]. Therefore, the ablated connectivity from E*β* population to any population *iα* is calculated as:

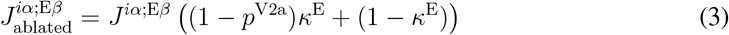

The inhibitory conductance strength after ablation of V1 or V2b neurons is calculated as:

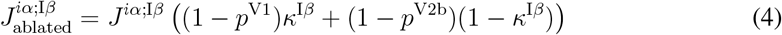

### Input currents and their modification with activation

The E and I neuronal populations receive external excitatory input, mimicking input from the brain or the dorsal spinal cord. This input enables the E neurons to fire and is modeled as an external depolarizing current 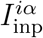. Activation of a population is modeled by injecting additional activation depolarizing current to that population 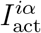. Therefore, the total input 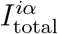 is

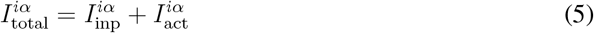

For I populations, when either V1 or V2b neurons are activated, the input current is calculated as:

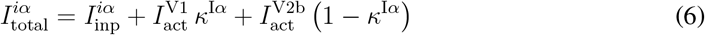

### Dynamics of neuronal populations

Each population is described by activation variables of the synaptic conductance *s*^*iα*^, firing rate *r*^*iα*^, adaptation current *a*^*iα*^ and depolarization current *d*^*iα*^. Synaptic facilitation is introduced into the model by setting the dynamics for facilitation variable *u*^*iα*^ [Tsodyks et al. (1998)]. The above-mentioned variables are described by the following equations:

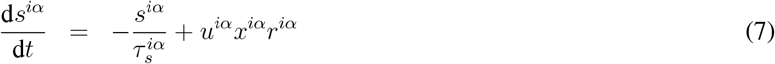

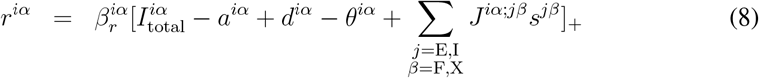

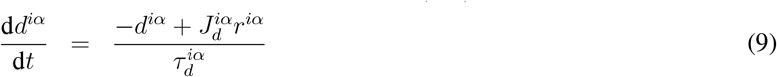

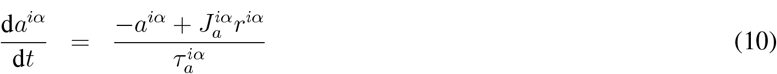

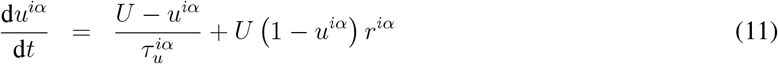

where 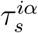 are synaptic time constants, 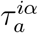 and 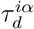 are the time constants of the adaptation and depolarization variables respectively, 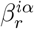 is slope of f-I curve (gain), 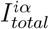 is total input current to a population (Eq. 5-6), *θ*^*iα*^ is threshold current, *J*^*iα*;*jβ*^are the synaptic conductance (Eq. 2-4), 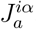 and 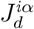 are ratio between *a*^*iα*^ or *d*^*iα*^ and *r*^*iα*^ at steady state, and *U* is the utilization of synaptic efficacy [Tsodyks et al. (1998)]. The Heavyside function [*x*]_+_ (Eq. 8) is defined as

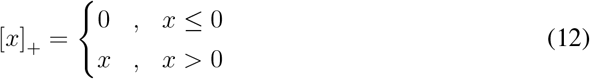

The threshold current *θ*^*iα*^ is incorporated into the default input current *I*_inp_, since they are both constant. Note that the time scale for dynamics of *a*^*iα*^ (*τ*_*a*_ *∼* 100 ms) is significantly larger than the time scale of *s*^*iα*^ (*τ*_*s*_ *∼* 1 ms). This makes adaptation current a slow current in our model, which is essential for generating bursting activities that were observed in experiments [Rinzel and Ermentrout (1998)]. The values *x*^*iα*^ are constants indicating available synaptic resource. We use the values *x*^E*α*^ = 0.92 for excitatory neurons and *x*^I*α*^ = *x*^M*α*^ = 0.48 for inhibitory neurons and motoneurons. Equation 11 is used only for the EF and EX populations. For the other populations, *u* = 0.4 is a fixed value.

### Parameters

The reference values of the parameter in our model are listed in Table 1, and are consistent with published datasets [Bikoff et al. (2016),Husch et al. (2015)].

### Single module model

To illustrate functions of neural circuit components, we simulated a circuit model made up of a single module. Parameters for neuronal populations and synaptic coupling coefficients in the single module models are the same as in the full circuit, except for some of the input currents and connectivity values that are modified to account for a smaller circuit. Parameters for the model that includes E and M populations (Figure S4B-D) are: 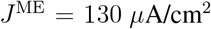 and 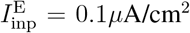. Parameters for the model that includes E, I and M populations (Figure 4C,D, S4E,F) aree, unless otherwise stated: *J* ^IE^ = 26 *µ*A/cm^2^, *J* ^EI^ = 10 *µ*A/cm^2^, *J* ^ME^ = 65 *µ*A/cm^2^, *J* ^MI^ = 100 *µ*A/cm^2^, and 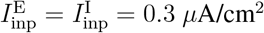.

## Biomechanical Model

To quantify the effects of spinal premotor interneuron manipulations on scratch frequency, we actuate a biomechanical model of the mouse ankle joint using motoneuron outputs from the neuronal model.

### Link Segment Model

We model the mouse ankle joint as a one degree-of-freedom, two-link segment model (Figure 4B). Only the distal segment is allowed to rotate around the axis of the fixed proximal segment. The parameters used in the biomechanical model are detailed in Table 2. The *y*-axis is parallel to the non-rotating segment, the *x*-axis points towards the right and the *z*-axis points towards the reader. The unit vectors along the axes are *ê*_*x*_, *ê*_*y*_ and *ê*_*z*_. The inertia of the distal segment around the axis of rotation is computed using the parallel axis theorem as:

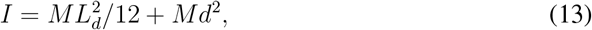

where *L*_*d*_ is the length of the distal segment, *M* is the mass of the segment, and *d* is the distance from the center of mass to the axis of rotation.

The distal segment is actuated by an antagonistic pair of identical muscles. The dynamics of the distal segment’s movement are described by Newton’s second law as:

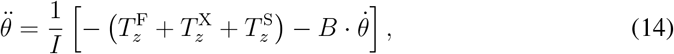

where 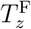 and 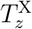 are the torques in the *z* direction generated by the flexor and extensor muscles respectively. The angle *θ* represents the ankle joint angle in radians, and 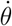 and 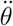 denote its first and second derivatives. *B* defines the external viscosity, which is set to 3 · 10^9^ dyn *×* cm *×* s. The fully extended ankle position is defined as *θ* = *π* with decreasing joint angles representing dorsiflexion, or ankle flexion. The torque from viscoelastic stops, *T*_S,z_, is used to constrain the range of motion using the following equation:

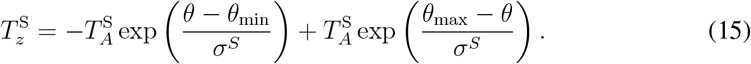

where 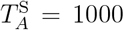 dyn and *σ*^S^ = 0.01 rad. The maximum joint angle (i.e. the maximum plantarflexion angle or extension), *θ*_max_, is set to 0.78 *π* (140^*°*^), and the minimum joint angle (i.e. the maximum flexion angle), *θ*_min_ to 0.39 *π* (70^*°*^).

The flexor and extensor muscles are connected in a bow-string configuration with the origin on the proximal segment *a* = 1.8 cm away from the rotation axis and the insertion on the distal segment *b* = 0.1 cm away from the rotation axis (Figure 4B) (Charles et al., 2016). The radius vector from the center of rotation *O* to the insertion point of the flexor muscle to the distal segment is

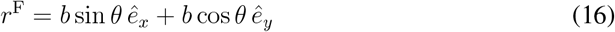

The length of the flexor muscle is

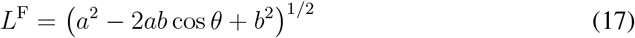

The vector force of the flexor muscle is

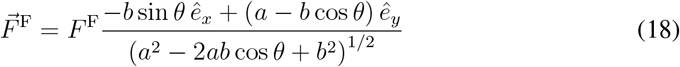

The torque of the flexor muscle is

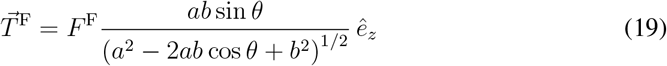

For the torque generated by the extensor muscle, a calculation similar to Eqs. 16-19 applies when *θ > θ*_*c*_ = *−* arccos (*b/a*), which corresponds to 1.63 rad (93.2^*°*^) for our parameter set. The length of the extensor muscle is calculated as

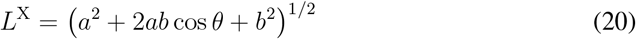

The torque of the extensor muscle as

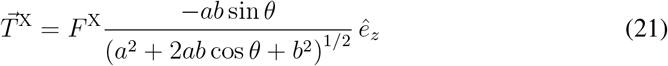

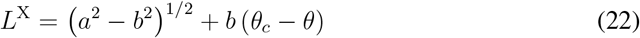

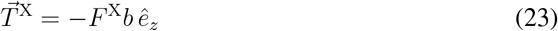

Note that “muscle length” here means the total length of the musculotendon unit, i.e. muscle fiber length 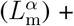 tendon/aponeurosis length 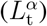 for *α* =F,X. For simplicity, we assume inflexible tendon/aponeurosis and therefore length changes are all taken up by muscle fibers. The optimal musculotendon length (the length of the musculotendon for which the active force is maximal) is taked to be *L*_0_ = 0.473 cm for both muscles (Charles et al., 2016). The maximal allowed length of the musculotendon is defined as *α*_*L*_*L*_0_ where *α*_*L*_ = 1.1. Therefore, the length of the tendon of the flexor is 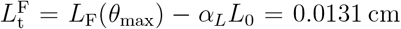 and that of the extensor is 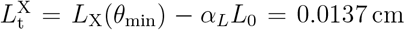 (Eq. 22). As the total muscle length *L*^*α*^(*t*) varies, the length of the musculotendon unit 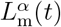 is

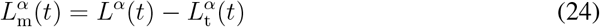

### Muscle Model

A muscle model converts motoneuron activity, *r*^MF^ and *r*^MX^, and muscle lengths into muscle forces, 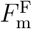 and 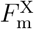 respectively. In addition to the low-pass filtering effects of muscle, we include the following known muscle properties: the length-dependent muscle activation, force-length relationship, and a passive elastic element. These properties are added to prevent drift in joint angles that would otherwise occur in a linear muscle model. Both flexor and extensor muscles are modeled identically to reduce the complexity of the model.

### Muscle force

The muscle force (*F*_m_) comprises forces from an active contractile element (*F*_CE_), and a passive elastic element (*F*_PE_) (Figure S7A). *F*_CE_ and *F*_PE_ are represented as a fraction of the muscle force, *F*_0_

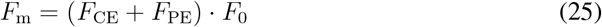

The maximum muscle force *F*_0_ is set to 2.13 · 10^5^ dyn for both muscles (Charles et al., 2016). *Passive elastic element*. The model includes a passive elastic element situated parallel to the contractile element. This passive elasticity arises from connective tissues within the muscle (Tsianos and Loeb, 2017). The force generated by this passive element, *F*_PE_, increases with the muscle fiber length (Figure S7B) (Song et al., 2008):

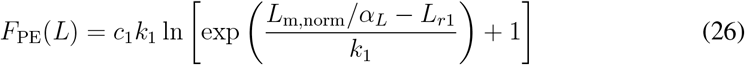

where the normalized musculotendon length is (Equation 24)

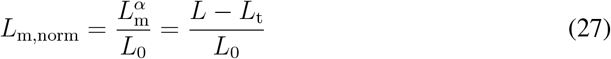

The parameters *k*_1_ and *L*_*r*1_ are obtained from (Song et al., 2008). The value of *c*_1_, which defines the asymptotic slope of the exponential function (i.e. stiffness), is adjusted to be 6 times larger than in the original model (Song et al., 2008) to account for the higher passive stiffness of the mouse muscle (Meyer and Lieber, 2018; Smith et al., 2011).

### Resting muscle length and joint angle

The resting joint angle *θ*_rest_ is the angle *θ* for which 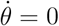 and 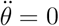 for zero active force. Assuming it is not near *θ*_min_ or *θ*_max_,

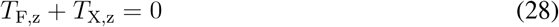

for *θ* = *θ*_rest_ and *F*_CE_ = 0 (Eq. 14). We solved Eqs. 28,19,21, 23,26 and obtained *θ*_*rest*_ = 1.31 rad (75^*°*^). This angle, together with 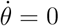, is used as initial condition for simulations.

### Active force

The amount of active force depends on the level of muscle activation, *A*, as well as the activation-dependent force-length relationship of the contractile element, *A*_*f*_ and *A*_FL_:

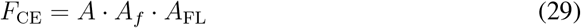

### Muscle activation

Neural excitation (i.e. the activity of motoneurons) results in the activation of cross-bridges within muscle fibers that can participate in active force generation, which we refer to as muscle activation *A*^*α*^(*t*); *α* =F,X. We used a model of second-order dynamics (Todorov, 2005) to relate *A*^*α*^(*t*) to the motoneuron firing rate *r*^M*α*^(*t*)

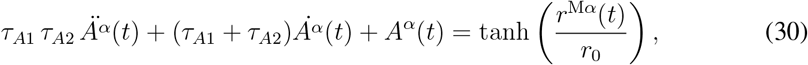

where *r*^M*α*^(*t*) is the population-average firing rate of the *α*th motoneuron pool (Eq. 8). and *r*_0_ = 54 Hz is a normalization constant. The tanh function in Equation 30 ensures that the normalized neural input to the muscle model is less than 1. *A*^*α*^(*t*) denotes muscle activation, which ranges from 0 to 1. This second-order differential equation captures the low-pass filtering effect of muscle (e.g. Baldissera et al. (1998)), arising from the biophysical properties of muscle force generation. *τ*_*A*1_ and *τ*_*A*2_ are set to 10 ms to account for the faster contraction capability of mouse muscles compared to humans and other animals (Martinez-Silva et al., 2018).

### Force-length relationship

The maximum amount of force a muscle can generate depends on its fiber length, known as the force-length relationship. This relationship scales muscle force as a function of the normalized muscle fiber length, *L*_m,norm_ (Eq. 27) (Song et al., 2008). The force-generating capability of a muscle decreases when the muscle becomes longer or shorter than its optimal muscle fiber length (*L*_m,norm_ = 1) (Rassier et al., 1999; Tsianos and Loeb, 2017). This property is thought to arise from the extent of myofilament overlap at different muscle lengths (Herzog et al., 1992). This nonlinear relationship is modeled by scaling muscle force by *A*_FL_ (Song et al., 2008):

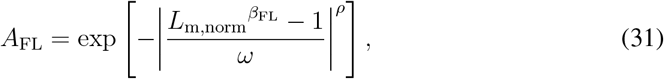

where the values of model parameters, *ω, β*_FL_, and *ρ* (Table 2) are those for fast-twitch units (Song et al., 2008).

We also include the dependence of muscle activation on muscle fiber length previously reported for mammalian skeletal muscles (Brown et al., 1999) (Figure S7C). This is thought to arise from mechanisms independent of myofilament overlap such as changes in the rate of cross-bridge formation and detachment (Brown et al., 1999). We model this by adding a scaling factor, *A*_*f*_, to muscle activation, whose value ranges from 0 to 1 (cf. Eq. 29).

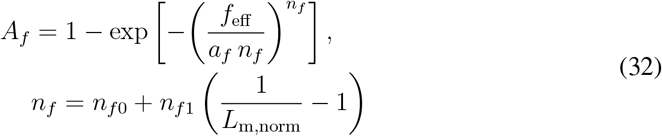

where

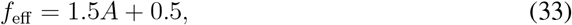

The parameters *a*_*f*_, *n*_*f*0_ and *n*_*f*1_, are taken from those for fast-twitch units in the original model (Brown et al., 1999). The total muscle-length curve is *A*_*f*_ · *A*_FL_ (Figure S7C)

## Calculation of oscillation frequency and identification of dynamics categories

### Frequency calculation

We use the function *find_peaks* in the python package *scipy*.*signal* to identify peaks in the simulated dynamics of MF and MX firing, as well as in the simulated joint angles. The number of peaks within a time interval *T*_interval_ is noted as *N*_peaks_ with a superscript “F”, “X” or “movement”, corresponding to peaks in *r*^MF^, *r*^MX^ and *θ* respectively. We defined the movement frequency as the number of joint movements varying more than *±*5^*°*^ from the balanced angle position (75^*°*^) per time unit, in sequences of “flexor-extensor” or “extensor-flexor” movements: 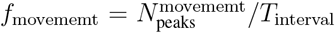. Therefore, if the movement does not reach 5^*°*^ in both directions, the joint is not considered to be oscillating (Figure 6F,6G).

A diversity of motoneuron dynamics emerge in our simulations. We divide the temporal dynamical states into four categories. In addition to regular, periodic firing, there are states of irregular oscillations, 100% co-contraction and atonia. Except during atonia, we compute the range of peak heights during a large time interval.

### Regular oscillations

(Figure 4G-J) are characterized by periodic oscillations with a non-zero phase difference between the MF and MX populations, and by the range of peak heights (difference between maximum and minimum peak heights) being less than 5% smaller than the mean.

### Irregular oscillations

(Figure S4J,K) are states in which the range of peak heights is 5% larger than the mean. The dynamics may be aperiodic.

In *100% co-contraction* states (Figure S4L,M), the MF and MX populations oscillate in phase and simultaneously reached their maximal value, leading to the joint angle being stuck or flexed or extended only in one direction. The dynamics are periodic and regular.

### Atonia

(Figure S4N,O) refers to a fixed point in the dynamical system, where its variables, including MF and MX, are constant with time.

### Co-contraction rate calculation

Besides 100% co-contraction states, partial co-contraction also happen in irregular oscillation states, during which some but not all MF/MX peaks oscillate in phase and simultaneously reach their maximal value (Figure S4J). To quantify the effects of ablation or activation of neuronal populations on the occurrence of co-contraction, we define co-contraction rate *r*^co-contract^ (Figure 6I) as

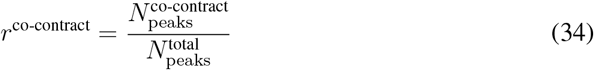

where 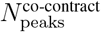 is the number of peaks in which MF and MX populations oscillate in phase and simultaneously reach their maximal values, and 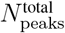 is the total number of peaks. During irregular oscillations, due to asymmetry, 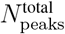 can be different for MF and MX, in such case we define 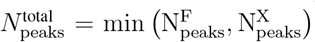. The value *r*^co-contract^ can be equivalently calculated from joint angle dynamics as 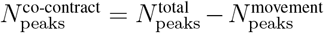. We categorize trials with r^co-contract^ *>* 0.99 as “100% co-contraction”.

## Simulations and visualization

### Numerical methods

Simulations of the neuro-mechanical model are performed using Euler’s method with time step Δ*t* = 0.1 ms.

### Simulations with random distributed connectivity values

For each tested parameter set, we repeated the simulations for 50 realizations with the values of synaptic conductance coefficients generated from a uniform distribution ranging between *±*10% of their assigned value (Table 1).

The computed values of a variable, such as oscillation frequency or phase difference between motoneuron populations, are plotted in graphs as mean *±* standard deviation (SD). There are cases in which there is almost no variability in the computed values despite variation in parameter set. For example, there is almost no variability in the phase difference between the flexor and extensor motorneuron populations (MF and MX) for V1 or V2b ablation efficiency above 0.30 (Figure S5F). This lack of variability is explained by the fact that bursting activity of one population occurs just after the activity of the other population. For example, the phase difference in Figure 4I (but not in Figure 4G) is dictated by the width of the burst.

### Initial conditions

For each realization of random connectivity values, the first simulation started with the following initial values: *a*^*iα*^(0) = 0.5 *µ*A*/*cm^2^, *d*^*iα*^(0) = 0 *µ*A*/*cm^2^, *u*^E*α*^(0) = 0.1, *s*^M*α*^(0) = *s*^EX^(0) = *s*^IX^(0) = 0.01, *s*^EF^(0) = 0.011, *s*^IF^(0) = 0.012. The initial values of *s*^*iα*^ are chosen to introduce a slight asymmetry between flexor and extensor at initialization, but the attractor dynamics are not sensitive to the choice of this asymmetry. Unless otherwise stated, trials following the first simulation are simulated using the final state of the last simulation as initial conditions.

### Computing the fractions of various dynamical categories

Each simulated trial is categorized as regular, irregular, 100% co-contraction or atonia (only in V1 or V2b activation case). The fraction of each dynamical category among the 50 realizations is computed. Only categories with fractions larger than 14% (7 out of 50 realizations) are plotted to ensure the generation of a sufficient amount of data to compute the statistics.

### Fourier analysis

To illustrate how the irregularity in oscillations changed following perturbations of population activity, Fourier spectrum of the joint angle dynamics is computed using *scipy*.*fft*.*rfft*. For each figure panel, a Fourier spectrum is computed for a randomly selected trial from the realizations.

## Notes

### Competing Interest Statement

The authors have declared no competing interest.

## Bibliography

Ampatzis, K., Song, J., Ausborn, J., and El Manira, A. (2014). Separate Microcircuit Modules of Distinct V2a Interneurons and Motoneurons Control the Speed of Locomotion. Neuron 83, 934– 943.

Ausborn, J., Mahmood, R., and El Manira, A. (2012). Decoding the rules of recruitment of excitatory interneurons in the adult zebrafish locomotor network. Proc. Natl. Acad. Sci. U. S. A. 109.

Ausborn, J., Snyder, A.C., Shevtsova, N.A., Rybak, I.A., and Rubin, J.E. (2018). State-dependent rhythmogenesis and frequency control in a half-center locomotor CPG. J. Neurophysiol. 119, 96– 117.

Ausborn, J., Shevtsova, N.A., and Danner, S.M. (2021). Computational Modeling of Spinal Locomotor Circuitry in the Age of Molecular Genetics. Int. J. Mol. Sci. 22.

Azim, E., Jiang, J., Alstermark, B., and Jessell, T.M. (2014). Skilled reaching relies on a V2a propriospinal internal copy circuit. Nature.

Baldissera, F., Cavallari, P., and Cerri, G. (1998). Motoneuronal pre-compensation for the lowpass filter characteristics of muscle. A quantitative appraisal in cat muscle units. J. Physiol. 511 (Pt 2), 611–627.

Bellardita, C., and Kiehn, O. (2015). Phenotypic characterization of speed-associated gait changes in mice reveals modular organization of locomotor networks. Curr. Biol. 25, 1426–1436.

Berg, R.W. (2017). Neuronal population activity in spinal motor circuits: Greater than the sum of its parts. Front. Neural Circuits 11, 294694.

Berg, R.W., Alaburda, A., and Hounsgaard, J. (2007). Balanced inhibition and excitation drive spike activity in spinal half-centers. Science 315, 390–393.

Berkowitz, A. (2010). Multifunctional and specialized spinal interneurons for turtle limb movements. Ann. N. Y. Acad. Sci. 1198, 119–132.

Berkowitz, A., and Hao, Z.-Z. (2011). Partly shared spinal cord networks for locomotion and scratching. Integr. Comp. Biol. 51, 890–902.

Berkowitz, A., Roberts, A., and Soffe, S.R. (2010). Roles for multifunctional and specialized spinal interneurons during motor pattern generation in tadpoles, zebrafish larvae, and turtles. Front. Behav. Neurosci. 4, 36.

Bikoff, J.B., Gabitto, M.I., Rivard, A.F., Drobac, E., Machado, T.A., Miri, A., Brenner-Morton, S., Famojure, E., Diaz, C., Alvarez, F.J., et al. (2016). Spinal Inhibitory Interneuron Diversity Delineates Variant Motor Microcircuits. Cell 165, 207–219.

Bourane, S., Duan, B., Koch, S.C., Dalet, A., Britz, O., Garcia-Campmany, L., Kim, E., Cheng, L., Ghosh, A., Ma, Q., et al. (2015). Gate control of mechanical itch by a subpopulation of spinal cord interneurons. Science (80-.). 350, 550–554.

Britz, O., Zhang, J., Grossmann, K.S., Dyck, J., Kim, J.C., Dymecki, S., Gosgnach, S., and Goulding, M. (2015). A genetically defined asymmetry underlies the inhibitory control of flexor-extensor locomotor movements. Elife 4.

Brown, T.G. (1911). The Intrinsic Factors in the Act of Progression in the Mammal. Proc. R. Soc. London B Biol. Sci. 84.

Brown, T.G. (1914). On the nature of the fundamental activity of the nervous centres; together with an analysis of the conditioning of rhythmic activity in progression, and a theory of the evolution of function in the nervous system. J. Physiol. 48, 18–46.

Brown, I.E., Cheng, E.J., and Loeb, G.E. (1999). Measured and modeled properties of mammalian skeletal muscle. II. The effects of stimulus frequency on force-length and force-velocity relationships. J. Muscle Res. Cell Motil. 20, 627–643.

Büschges, A., Akay, T., Gabriel, J.P., and Schmidt, J. (2008). Organizing network action for locomotion: insights from studying insect walking. Brain Res. Rev. 57, 162–171.

Caldeira, V., Dougherty, K.J., Borgius, L., and Kiehn, O. (2017). Spinal Hb9::Cre-derived excitatory interneurons contribute to rhythm generation in the mouse. Sci. Reports 2017 71 7, 1– 12.

Callahan, R.A., Roberts, R., Sengupta, M., Kimura, Y., Higashijima, S.I., and Bagnall, M.W. (2019). Spinal V2b neurons reveal a role for ipsilateral inhibition in speed control. Elife 8.

Charles, J.P., Cappellari, O., Spence, A.J., Wells, D.J., and Hutchinson, J.R. (2016). Muscle moment arms and sensitivity analysis of a mouse hindlimb musculoskeletal model. J. Anat. 229, 514–535.

Coombes, S., and Bressloff, P.C. (2005). Bursting: The genesis of rhythm in the nervous system (World Scientific Publishing Co.).

Crone, S.A., Quinlan, K.A., Zagoraiou, L., Droho, S., Restrepo, C.E., Lundfald, L., Endo, T., Setlak, J., Jessell, T.M., Kiehn, O., et al. (2008). Genetic ablation of V2a ipsilateral interneurons disrupts left-right locomotor coordination in mammalian spinal cord. Neuron 60, 70–83.

Crone, S.A., Zhong, G., Harris-Warrick, R., and Sharma, K. (2009). In mice lacking V2a interneurons, gait depends on speed of locomotion. J Neurosci 29, 7098–7109.

Deliagina, T.G., Feldman, A.G., Gelfand, I.M., and Orlovsky, G.N. (1975). On the role of central program and afferent inflow in the control of scratching movements in the cat. Brain Res. 100, 297–313.

Dougherty, K.J., and Ha, N.T. (2019). The rhythm section: an update on spinal interneurons setting the beat for mammalian locomotion. Curr. Opin. Physiol. 8, 84–93.

Dougherty, K.J., and Kiehn, O. (2010). Firing and cellular properties of V2a interneurons in the rodent spinal cord. J. Neurosci. 30, 24–37.

Dougherty, K.J., Zagoraiou, L., Satoh, D., Rozani, I., Doobar, S., Arber, S., Jessell, T.M., and Kiehn, O. (2013). Locomotor rhythm generation linked to the output of spinal shox2 excitatory interneurons. Neuron 80, 920–933.

Friedman, A.K., Zhurov, Y., Ludwar, B.C., and Weiss, K.R. (2009). Motor outputs in a multitasking network: relative contributions of inputs and experience-dependent network states. J. Neurophysiol. 102, 3711–3727.

Friesen, W.O., and Stent, G.S. (1977). Generation of a locomotory rhythm by a neural network with recurrent cyclic inhibition. Biol. Cybern. 28, 27–40.

Frigon, A. (2012). Central pattern generators of the mammalian spinal cord. Neuroscientist 18, 56–69.

Frigon, A., and Gossard, J.P. (2010). Evidence for specialized rhythm-generating mechanisms in the adult mammalian spinal cord. J. Neurosci. 30, 7061–7071.

Golomb, D., Shedmi, A., Curtu, R., and Ermentrout, G.B. (2006). Persistent synchronized bursting activity in cortical tissues with low magnesium concentration: a modeling study. J. Neurophysiol. 95, 1049–1067.

Golomb, D., Moore, J.D., Fassihi, A., Takatoh, J., Prevosto, V., Wang, F., and Kleinfeld, D. (2022). Theory of hierarchically organized neuronal oscillator dynamics that mediate rodent rhythmic whisking. Neuron 110, 3833-3851.e22.

Gosgnach, S., Lanuza, G.M., Butt, S.J., Saueressig, H., Zhang, Y., Velasquez, T., Riethmacher, D., Callaway, E.M., Kiehn, O., and Goulding, M. (2006). V1 spinal neurons regulate the speed of vertebrate locomotor outputs. Nature 440, 215–219.

Goulding, M. (2009). Circuits controlling vertebrate locomotion: moving in a new direction. Nat Rev Neurosci 10, 507–518.

Hayashi, M., Hinckley, C.A., Driscoll, S.P., Moore, N.J., Levine, A.J., Hilde, K.L., Sharma, K., and Pfaff, S.L. (2018). Graded Arrays of Spinal and Supraspinal V2a Interneuron Subtypes Underlie Forelimb and Hindlimb Motor Control. Neuron 97.

Hayashi, M., Gullo, M., Senturk, G., Costanzo S. Di, Nagasaki, S.C., Kageyama, R., Imayoshi, I., Goulding, M., Pfaff, S.L., and Gatto, G. (2023). A spinal synergy of excitatory and inhibitory neurons coordinates ipsilateral body movements. Elife 12.

Hayut, I., Fanselow, E.E., Connors, B.W., and Golomb, D. (2011). LTS and FS inhibitory interneurons, short-term synaptic plasticity, and cortical circuit dynamics. PLoS Comput. Biol. 7.

Herzog, W., Kamal, S., and Clarke, H.D. (1992). Myofilament lengths of cat skeletal muscle: theoretical considerations and functional implications. J. Biomech. 25, 945–948.

Husch, A., Dietz, S.B., Hong, D.N., and Harris-Warrick, R.M. (2015). Adult spinal V2a interneurons show increased excitability and serotonin-dependent bistability. J. Neurophysiol. 113, 1124–1134.

Jackman, S.L., and Regehr, W.G. (2017). The Mechanisms and Functions of Synaptic Facilitation. Neuron 94, 447–464.

Jankowska, E., Jukes, M.G.M., Lund, S., and Lundberg, A. (1967a). The effect of DOPA on the spinal cord. 5. Reciprocal organization of pathways transmitting excitatory action to alpha motoneurones of flexors and extensors. Acta Physiol. Scand. 70, 369–388.

Jankowska, E., Jukes, M.G.M., Lund, S., and Lundberg, A. (1967b). The Effect of DOPA on the Spinal Cord 6. Half-centre organization of interneurones transmitting effects from the flexor reflex afferents. Acta Physiol. Scand. 70, 389–402.

Kiehn, O. (2016). Decoding the organization of spinal circuits that control locomotion. Nat. Rev. Neurosci. 17, 224–238.

Kimura, Y., and Higashijima, S. ichi (2019). Regulation of locomotor speed and selection of active sets of neurons by V1 neurons. Nat. Commun. 2019 101 10, 1–12.

Kishore, S., Bagnall, M.W., and McLean, D.L. (2014). Systematic Shifts in the Balance of Excitation and Inhibition Coordinate the Activity of Axial Motor Pools at Different Speeds of Locomotion. J. Neurosci. 34, 14046–14054.

Lindén, H., and Berg, R.W. (2021). Why Firing Rate Distributions Are Important for Understanding Spinal Central Pattern Generators. Front. Hum. Neurosci. 15, 719388.

Lindén, H., Petersen, P.C., Vestergaard, M., and Berg, R.W. (2022). Movement is governed by rotational neural dynamics in spinal motor networks. Nature 610, 526–531.

Lundberg, A. (1981). Half-centres revisited. In Regulatory Functions of the CNS. Motion and Organization Principles, J. Szentagotheu, M. Palkovits, and Hamori J, eds. (Pergamon Akademiai Kiado), pp. 155–167.

Marder, E., and Bucher, D. (2001). Central pattern generators and the control of rhythmic movements. Curr. Biol. 11, R986–R996.

Marder, E., and Calabrese, R.L. (1996). Principles of rhythmic motor pattern generation. Physiol. Rev. 76, 687–717.

Martínez-Silva, M. de L., Imhoff-Manuel, R.D., Sharma, A., Heckman, C.J., Shneider, N.A., Roselli, F., Zytnicki, D., and Manuel, M. (2018). Hypoexcitability precedes denervation in the large fast-contracting motor units in two unrelated mouse models of ALS. Elife 7.

McCrea, D.A., and Rybak, I.A. (2008). Organization of mammalian locomotor rhythm and pattern generation. Brain Res. Rev. 57, 134–146.

Melamed, O., Barak, O., Silberberg, G., Markram, H., and Tsodyks, M. (2008). Slow oscillations in neural networks with facilitating synapses. J. Comput. Neurosci. 25, 308–316.

Meyer, G., and Lieber, R.L. (2018). Muscle fibers bear a larger fraction of passive muscle tension in frogs compared with mice. J. Exp. Biol. 221.

Pehlevan, C., and Sompolinsky, H. (2014). Selectivity and sparseness in randomly connected balanced networks. PLoS One 9.

Perkel, D.H., and Mulloney, B. (1974). Motor Pattern Production in Reciprocally Inhibitory Neurons Exhibiting Postinhibitory Rebound. Science (80-.). 185, 181–183.

Pinto, C.M.A., and Golubitsky, M. (2006). Central pattern generators for bipedal locomotion. J. Math. Biol. 53, 474–489.

Rassier, D.E., MacIntosh, B.R., and Herzog, W. (1999). Length dependence of active force production in skeletal muscle. J. Appl. Physiol. 86, 1445–1457.

Reid, B., Slater, C.R., and Bewick, G.S. (1999). Synaptic vesicle dynamics in rat fast and slow motor nerve terminals. J. Neurosci. 19, 2511–2521.

Rinzel, J., and Ermentrout, G.B. (1998). Analysis of neural excitability and oscillations. Methods Neuronal Model. 251–292.

Rybak, I.A., Shevtsova, N.A., Lafreniere-Roula, M., and McCrea, D.A. (2006). Modelling spinal circuitry involved in locomotor pattern generation: insights from deletions during fictive locomotion. J. Physiol. 577, 617–639.

Schuster, H.G., and Just, W. (2005). Deterministic Chaos: An Introduction: Fourth Edition. Determ. Chaos An Introd. Fourth Ed. 1–287.

Sciolino, N.R., Plummer, N.W., Chen, Y.W., Alexander, G.M., Robertson, S.D., Dudek, S.M., McElligott, Z.A., and Jensen, P. (2016). Recombinase-Dependent Mouse Lines for Chemogenetic Activation of Genetically Defined Cell Types. Cell Rep. 15, 2563–2573.

Sengupta, M., and Bagnall, M.W. (2023). Spinal Interneurons: Diversity and Connectivity in Motor Control. Annu. Rev. Neurosci. 46, 79–99.

Sherrington, C.S. (1910). NOTES ON THE SCRATCH-REFLEX OF THE CAT. Q. J. Exp. Physiol. 3, 213–220.

Sherwood, W.E., Harris-Warrick, R., and Guckenheimer, J. (2011). Synaptic patterning of left-right alternation in a computational model of the rodent hindlimb central pattern generator. J. Comput. Neurosci. 30, 323–360.

Shriki, O., Hansel, D., and Sompolinsky, H. (2003). Rate models for conductance-based cortical neuronal networks. Neural Comput. 15, 1809–1841.

Skinner, F.K., Kopell, N., and Marder, E. (1994). Mechanisms for oscillation and frequency control in reciprocally inhibitory model neural networks. J. Comput. Neurosci. 1, 69–87.

Smith, L.R., Lee, K.S., Ward, S.R., Chambers, H.G., and Lieber, R.L. (2011). Hamstring contractures in children with spastic cerebral palsy result from a stiffer extracellular matrix and increased in vivo sarcomere length. J. Physiol. 589, 2625–2639.

Soloduchin, S., and Shamir, M. (2018). Rhythmogenesis evolves as a consequence of long-term plasticity of inhibitory synapses. Sci. Rep. 8.

Song, D., Raphael, G., Lan, N., and Loeb, G.E. (2008). Computationally efficient models of neuromuscular recruitment and mechanics. J. Neural Eng. 5, 175.

Song, J., Dahlberg, E., and El Manira, A. (2018). V2a interneuron diversity tailors spinal circuit organization to control the vigor of locomotor movements. Nat. Commun. 9.

Song, J., Pallucchi, I., Ausborn, J., Ampatzis, K., Bertuzzi, M., Fontanel, P., Picton, L.D., and El Manira, A. (2020). Multiple Rhythm-Generating Circuits Act in Tandem with Pacemaker Properties to Control the Start and Speed of Locomotion. Neuron 105, 1048-1061.e4.

Strohmer, B., Najarro, E., Ausborn, J., Berg, R.W., and Tolu, S. (2024). Sparse Firing in a Hybrid Central Pattern Generator for Spinal Motor Circuits. Neural Comput. 36, 759–780.

Talpalar, A.E., Bouvier, J., Borgius, L., Fortin, G., Pierani, A., and Kiehn, O. (2013). Dual-mode operation of neuronal networks involved in left–right alternation. Nature 500, 85–88.

Todorov, E. (2005). Stochastic optimal control and estimation methods adapted to the noise characteristics of the sensorimotor system. Neural Comput. 17, 1084–1108.

Trevisan, A.J., Han, K., Chapman, P., Kulkarni, A.S., Hinton, J.M., Ramirez, C., Klein, I., Gatto, G., Gabitto, M.I., Menon, V., et al. (2024). The transcriptomic landscape of spinal V1 interneurons reveals a role for En1 in specific elements of motor output. BioRxiv 2024.09.18.613279.

Tsianos, G.A., and Loeb, G.E. (2017). Muscle and Limb Mechanics. Compr. Physiol. 7, 429–462.

Tsodyks, M., Pawelzik, K., and Markram, H. (1998). Neural networks with dynamic synapses. Neural Comput. 10, 821–835.

Vijatovic, D., Toma, F.A., Harrington, Z.P.M., Sommer, C., Hauschild, R., Trevisan, A.J., Chapman, P., Julseth, M.J., Brenner-Morton, S., Gabitto, M.I., et al. (2024). Spinal neuron diversity scales exponentially with swim-to-limb transformation during frog metamorphosis. BioRxiv 2024.09.20.614050.

Van Vreeswijk, C., and Hansel, D. (2001). Patterns of synchrony in neural networks with spike adaptation. Neural Comput. 13, 959–992.

Van Vreeswijk, C., and Sompolinsky, H. (1996). Chaos in neuronal networks with balanced excitatory and inhibitory activity. Science (80-.). 274, 1724–1726.

Wallén, P., and Grillner, S. (1987). N-methyl-D-aspartate receptor-induced, inherent oscillatory activity in neurons active during fictive locomotion in the lamprey. J. Neurosci. 7, 2745–2755.

Zhang, H., Shevtsova, N.A., Deska-Gauthier, D., Mackay, C., Dougherty, K.J., Danner, S.M., Zhang, Y., and Rybak, I.A. (2022). The role of V3 neurons in speeddependent interlimb coordination during locomotion in mice. Elife 11, 2021.09.01.458603.

Zhang, J., Lanuza, G.M., Britz, O., Wang, Z., Siembab, V.C., Zhang, Y., Velasquez, T., Alvarez, F.J., Frank, E., and Goulding, M. (2014). V1 and v2b interneurons secure the alternating flexor-extensor motor activity mice require for limbed locomotion. Neuron 82, 138–150.

Zhang, Y., Narayan, S., Geiman, E., Lanuza, G.M., Velasquez, T., Shanks, B., Akay, T., Dyck, J., Pearson, K., Gosgnach, S., et al. (2008). V3 spinal neurons establish a robust and balanced locomotor rhythm during walking. Neuron 60, 84–96.

Zhong, G., Droho, S., Crone, S.A., Dietz, S., Kwan, A.C., Webb, W.W., Sharma, K., and Harris-Warrick, R.M. (2010). Electrophysiological Characterization of V2a Interneurons and Their Locomotor-Related Activity in the Neonatal Mouse Spinal Cord. J. Neurosci. 30, 170–182.

