## Supplemental Figure Legend for "The spinal premotor network driving scratching flexor and extensor alternation"

**Figure S1. High-frequency flexor and extensor oscillations during scratching (refers to Figure 1).**

A,B) Line graphs representing the height (y-coordinates) of knee (green), ankle (dark orange), and hindpaw (light orange) during scratching in control mice. Grey shaded areas label individual scratch episodes, showing the variability in oscillation numbers and duration, and final position of the paw at the end of individual episodes.

C) Bar graph showing the duration of the approach phase, reconstructed from high-speed videos, in 3 control mice. Individual mice are represented as grey dots.

D,E) Line graphs representing the height of knee (green), ankle (dark orange), and hindpaw (light orange) during scratching episodes of high (D, ~20 Hz) or moderate (E, ~10 Hz) frequency.

**Figure S2. Ipsilateral excitatory V2a neurons drive the high-frequency scratch oscillations (refers to Figure 2).**

A,C,G) Schematics illustrating the ablation of (A), the CNO-driven silencing (C), and the CNO-driven activation (G) of V2a neurons.

B) Bar graph showing the reduction in number of scratch bouts in V2a neuron-ablated mice compared to controls, $p = 0.001$.

D,E) Bar graphs showing unchanged number of scratch bouts, $p = 0.2408$ (D), and reduced oscillation frequency, $p < 0.0001$ (E), in V2a neuron-silenced mice compared to controls.

F) Frequency distributions of the speed of oscillations (bouts/second) of all episodes occurring in the 30 minutes of recorded scratch response in control (red, N=7) and V2a neuron-silenced (yellow, N=5) mice. SEM represented as shaded area. Statistical analysis performed using two-way ANOVA (interaction genotype x speed, $p < 0.0001$) with Bonferroni's *post-hoc* test, $p < 0.0001$ for genotype comparison at 2, 4, and 6 bouts/sec.

H,I) Bar graphs showing similar number of scratch bouts, $p = 0.2546$ (H) and increase in frequency of oscillations, $p = 0.0001$ (I), in mice with CNO-activated V2a neurons compared to controls.

Data presented as mean ± SEM. Individual mice represented as filled grey circles. Statistical analysis performed using two-tailed Student's *t-test*, unless otherwise indicated.

**Figure S3. Ipsilateral inhibitory neurons modulate the frequency of scratch oscillations (refers to Figure 3).**

A,C,D) Schematics illustrating the ablation of V1 neurons (A), the CNO-driven silencing of V1 neurons (C), and the ablation of V2b neurons (D).

B) Bar graph showing the reduction in number of scratch bouts in V1 neuron-ablated mice compared to controls, $p = 0.0371$.

D,E) Bar graphs showing the unchanged number of scratch bouts, $p = 0.1373$ (D), and the reduction in oscillation frequency, $p = 0.0112$ (E), in V1 neuron-silenced mice compared to controls.

F) Frequency distributions of the speed of oscillations (bouts/second) of all episodes occurring in the 30 minutes of recorded scratch response in control (dark cerulean, N=5) and V1 neuron-silenced (light violet, N=7) mice. Statistical analysis performed using two-way ANOVA (interaction genotype x speed, $p = 0.0034$) with Bonferroni's *post-hoc* test, for genotype comparison $p = 0.0039$ at 2 bouts/sec, $p = 0.0238$ at 6 bouts/sec.

H) Bar graph showing the unchanged number of scratch bouts in V2b neuron-ablated mice compared to controls, $p = 0.7560$.

I) Frequency distribution of the speed of oscillations (bouts/second) of all episodes occurring in the 30 minutes of recorded scratch responses in control (midnight blue, N=3) and V2b neuron-ablated (navy blue, N=4) mice. Statistical analysis performed using two-way ANOVA (interaction genotype x speed, $p = 0.0005$) with Bonferroni's *post-hoc* test, $p = 0.0346$ at 2 bouts/sec, $p = 0.0084$ at 4 bouts/sec, $p = 0.0063$ at 8 bouts/sec for genotype comparison.

Data presented as mean ± SEM, SEM represented as shaded area in panels F and I. Individual mice represented as filled grey circles. Statistical analysis performed using two-tailed Student's *t-test*, unless otherwise indicated.

**Figure S4. Neuromechanical model for high-frequency flexor and extensor alternation. (refers to Figure 4).**

A) Bar graph showing the asymmetry in number of V1 and V2b neurons contributing to the IF and IX populations.

B) Schematic illustrating the rhythm generator consisting of a single excitatory (E) population driving motoneuron (M) activity.

C,D) Simulated time traces of motoneuron firing rates (C) and E neuron intrinsic adaptation and depolarization currents (D). These two currents counteract each other, thus contributing to generating rhythmic bursting oscillations.

E) Left: Schematic illustrating a single module including coupled excitatory (E) and inhibitory (I) neurons and a motoneuron (M) population driven by E. Right: Simulated time traces of the intrinsic adaptation and intrinsic depolarization currents for three distinct strengths of I-to-E inhibitory conductance $J^{EI}$.

F) Simulated time traces of motoneuron firing rates in a single module circuit for three distinct strengths of E-to-I excitatory conductance $J^{IE}$.

G) Simulations of the oscillation frequency generated by the neuromechanical model as a function of the strength of the inter-module I-to-E inhibition ($J^{\mathrm{EI}}_{\mathrm{inter}}$). Red arrowhead and number indicate the value of synaptic strength used as reference parameter in our rate model (**Table 1**).

H, I) Graphs showing how the dependence of the computed oscillation frequency on the ablation efficiency ($p$) of inhibitory neurons (no discrimination between V1 and V2b neurons) changes for distinct values of inter-module E-to-I inhibition ($J^{\mathrm{EI}}_{\mathrm{inter}}$) (H) and I-to-I inhibition ($J^{\mathrm{II}}_{\mathrm{inter}}$) (I).

J-M) Simulated firing rates of MF and MX (J,L) and joint angle θ (K,M) generated by the neuromechanical model when 80% ($p^{\mathrm{V1}}$=0.8) (J,K) and 95% ($p^{\mathrm{V1}} = p^{\mathrm{V2a}}$ =0.95) (L,M) of V1 neuron-driven inhibition is eliminated. Red arrowheads denote co-contraction events.

N,O) Simulated firing rates of MF and MX (N) and joint angle θ (O) generated by injecting a current ($I^{\mathrm{V1}}_{\mathrm{act}}$) of 2 μA/cm$^2$ into V1 neurons.

Modeling data presented as mean ± SD of 50 realizations of synaptic conductance parameters, SD represented as shaded area.

**Figure S5. Computed circuits dynamics induced by manipulating the activity of individual neuronal populations (refers to Figure 5).**

A-C) Fourier transform of simulated ankle joint traces for increasing ablation efficiency of V2a (A), V1 (B), orV2b (C) neurons.

D) Graphs showing the computed oscillation frequency as a function of the inter-module E-to-I excitation ($J^{\mathrm{IE}}_{\mathrm{inter}}$) and for several values of the intra-module I-to-E inhibition ($J^{\mathrm{EI}}_{\mathrm{intra}}$).

E,F) Phase difference between the peaks of MF and MX activity as function of V1 or V2b neuron ablation efficiency, computed for one (E) and for 50 realizations (F). Only stable periodic MF and MX oscillations were used to calculate phase differences.

G) Fourier transform of simulated ankle joint oscillations for increasing values of current injected into V2a neurons.

H) Graph showing the oscillation frequency as a function of the addition to input current to E neurons due to their chemogenetic activation with facilitation ($u^{\mathrm{FX}}$ and $u^{\mathrm{EX}}$ vary according to

Equation 11) and without facilitation ($u^{EX} = u^{EX} = U^{E}$, **Table 1**).

I,J) Graph showing the oscillation frequency as a function of the additional input current for distinct values of the utilization of synaptic efficacy, $U^{E}$ (I) and of the facilitation time constant, $\tau_{u}^{E}$ (J).

K,L) Fourier transform of simulated ankle joint oscillations for increasing values of current injected in V1 (K) and V2b (L) neurons.

Modeling data presented as mean ± SD of 50 realizations of synaptic conductance parameters, SD represented as shaded area. The spectral power of each trace in A-C, G, K, L is normalized to a maximum value of 1.0.

**Figure S6. Distinct cooperation dynamics among the ipsilateral neuron populations in driving oscillations frequency (refers to Figure 6).**

A,E,I,M) Schematics illustrating the ablation of V2a and V2b neurons (A), the ablation of V1 and V2a neurons (E), the CNO-driven activation of V1 and V2a neurons (I), and the activation of V2a and V2b neurons (M).

B) Bar graph showing the reduction in number of scratch episodes in V2a neuron-ablated, V2b neuron-ablated, and dual V2a and V2b neuron-ablated mice compared to littermate controls. Statistical analysis was performed using two-tailed Student's *t-test*, $p = 0.0066$ V2a and V2b control vs ablated, $p = 0.0713$ V2b vs V2a ablated, $p = 0.3733$ V2b vs V2a and V2b ablated, $p = 0.1073$ V2a ablated vs V2a and V2b ablated.

C) Bar graph showing the computed reductions in oscillation frequency following ablation of V2a, V2b, and both V2a and V2b neurons in the neuromechanical model. Ablation efficiency is $p^{V2b} = 0.95$ for V2b ablation, $p^{V2a} = 0.95$ for V2a ablation, and $p^{V2a} = p^{V2b} = 0.95$ for both V2a and V2b ablation.

D) Fourier transform of simulated ankle joint oscillations for increasing ablation efficiency (*p*) of V2a and V2b neurons.

F) Bar graph showing the reduction in number of scratch episodes in V1 neuron-ablated, V2a neuron-ablated, and dual V1 and V2a neuron-ablated mice compared to littermate controls, $p < 0.0001$ V1 and V2a control vs ablated, $p = 0.1043$ V1 vs V2a ablated, $p = 0.020$ V1 vs V1 and V2a ablated, $p = 0.0003$ V2a ablated vs V1 and V2a ablated.

G) Bar graph showing the computed reductions in oscillation frequency as function of the ablation efficiency of V1, V2a, and both V1 and V2a neurons in the neuromechanical model. Ablation efficiency is $p^{V1} = 0.95$ for V1 ablation, $p^{V2a} = 0.95$ for V2a ablation, and $p^{V2a} = p^{V1} = 0.95$ for both V2a and V1 ablation.

H) Fourier transform of simulated ankle joint oscillations for increasing ablation efficiency ($p$) of V1 and V2a neurons.

J) Bar graph showing the decreased number of scratch episodes in V1 neuron-activated mice, and the lack of changes following activation of V2a and dual V1 and V2a neurons, $p = 0.9646$ V1 and V2a control vs activated, $p = 0.0039$ V1 vs V2a activated, $p = 0.0025$ V1 vs V1 and V2a activated, $p = 0.1512$ V2a activated vs V1 and V2a activated.

K) Bar graph showing computed increases in oscillation frequency as function of the activation of V1, V2a, and both V1 and V2a neurons in the neuromechanical model. Additional input current is $I_{act}^{V1} = 5\mu A/cm^2$ for V1 activation, $I_{act}^{V2a} = 5\mu A/cm^2$ for V2a activation, and $I_{act}^{V2a} = I_{act}^{V1} = 5\mu A/cm^2$ for both V1 and V2a activation.

L) Fourier transform of simulated ankle joint oscillations for increasing values of injected current ($I_{act}$) in V1 and V2a neurons.

N) Graphs showing the computed oscillation frequency and type of motoneuron firing as function of the amount of current injected ($I_{act}$) in V2a, V2b, and both V2a and V2b neurons. The bars above represent the occurrence of distinct motoneuron firing patterns (e.g., regular, irregular) as amount of injected current increases.

O) Fourier transform of simulated ankle joint oscillations for increasing values of injected current ($I_{act}$) in V2a and V2b neurons.

Experimental data presented as mean ± SEM, individual mice represented as filled grey circles. Statistical analysis performed using two-tailed Student's *t-test.* Modeling data presented as mean ± SD of 50 realizations, SD represented as shaded area in N. In panels D, H, L, O the spectral power of each trace is normalized to a maximum value of 1.0.

**Figure S7. Data-driven neuromechanical model of ipsilateral rhythmic oscillations (refers to Figure 4).**

A) Schematic illustrating that the total muscle force ($F_m$) includes forces from an active contractile element ($F_{CE}F_0$), and a passive elastic element ($F_{PE}F_0$).

B) Dependence of normalized passive muscle force $F_{PE}$ on the normalized muscle length $L_{m,norm}$. The passive muscle force increases approximately exponentially with increasing muscle length up to its anatomical maximum ($L_{m,norm} = \alpha_L$).

C) Dependence of total force-length curve on the normalized muscle length $L_{m,norm}$ and for several values of the variable $f_{eff}$ that depends on the muscle activation $A$. For the larger values of $f_{eff}$, the total force-length curve peaks near the optimal muscle length ($L_{m,norm} = 1$). It peaks at larger values of the length $L_{m,norm}$ for smaller $f_{eff}$.
