## Supplemental Table 1 for "The spinal premotor network driving scratching flexor and extensor alternation"

**Table 1: Model parameters for the neuronal model.**

| Parameter definition | Symbol | Value |
|---|---|---|
| Input current | $I_{\mathrm{inp}}^{\mathrm{E}\alpha}$ | 0.6 μA/cm$^2$ |
| | $I_{\mathrm{inp}}^{\mathrm{I}\alpha}$ | 0.3 μA/cm$^2$ |
| | $I_{\mathrm{inp}}^{\mathrm{M}\alpha}$ | 0 |
| Fraction of V1 neurons in I populations | $\kappa^{\mathrm{IX}}$ | 0.82 |
| | $\kappa^{\mathrm{IF}}$ | 0.45 |
| Synaptic time constant | $\tau_{\mathrm{s}}^{\mathrm{E}\alpha}$ | 4.0 ms |
| | $\tau_{\mathrm{s}}^{\mathrm{I}\alpha}$, $\tau_{\mathrm{s}}^{\mathrm{M}\alpha}$ | 5.4 ms |
| Facilitation time constant | $\tau_{\mathrm{u}}^{\mathrm{E}\alpha}$ | 250 ms |
| Adaptation current time constant | $\tau_{\alpha}^{\mathrm{E}\alpha}$ | 167 ms |
| | $\tau_{\alpha}^{\mathrm{I}\alpha}$ | 63 ms |
| | $\tau_{\alpha}^{M\alpha}$ | 500 ms |
| Adaptation current strength | $J_{\alpha}^{\mathrm{E}\alpha}$ , $J_{\alpha}^{\mathrm{M}\alpha}$ | 140 ms × μA/cm$^2$ |
| | $J_{\alpha}^{\mathrm{I}\alpha}$ | 180 ms × μA/cm$^2$ |
| Depolarization current time constant | $\tau_{d}^{\mathrm{E}\alpha}$ | 1.9 ms |
| Depolarization current strength | $J_{d}^{\mathrm{E}\alpha}$ | 110 ms × μA/cm$^2$ |
| Slope of f-I curve | $\beta_{\mathrm{r}}$ | 0.011 cm$^2$/(ms × μA) |
| Utilization of synaptic efficacy | $U^{\mathrm{E}\alpha}$ | 0.03 |
| | $u^{\mathrm{I}\alpha}, u^{\mathrm{M}\alpha}$ | 0.4 |
| Available synaptic resource | $x^{\mathrm{E}\alpha}$ | 0.92 |
| | $x^{\mathrm{I}\alpha}, x^{\mathrm{M}\alpha}$ | 0.48 |
| Synaptic coupling coefficients | $J_{\mathrm{intra}}^{\mathrm{IE}}$ | 52 μA/cm$^2$ |
| | $J_{\mathrm{intra}}^{\mathrm{ME}}$ | 195 μA/cm$^2$ |
| | $J_{\mathrm{inter}}^{\mathrm{EE}}$ | 6.5 μA/cm$^2$ |
| | $J_{\mathrm{intra}}^{\mathrm{EI}}$ | 66 μA/cm$^2$ |
| | $J_{\mathrm{inter}}^{\mathrm{EI}}$ | 77 μA/cm$^2$ |
| | $J_{\mathrm{inter}}^{\mathrm{II}}$ | 55 μA/cm$^2$ |
| | $J_{\mathrm{inter}}^{\mathrm{MI}}$ | 330 μA/cm$^2$ |

When an index α is written, the parameter value is valid for α = F, X.
Synaptic coupling coefficients that are not listed in this table are 0.
