## Supplemental Table 2 for "The spinal premotor network driving scratching flexor and extensor alternation"

**Table 2: Model parameters for the biomechanical model**

| Parameter definition | Symbol | Value |
|---|---|---|
| Segment mass | $M$ | 0.4 g |
| Segment length | $L_d$ | 1 cm |
| Distance between rotation axis and segment center of mass | $d$ | 0.1 cm |
| Distance between muscle contact point and insertion point | a | 1.8 cm |
| Distance between rotation axis and muscle contact point | b | 0.1 cm |
| Joint viscosity | B | $3 \times 10^9$ dyn × cm × s |
| Optimal muscle length | $L_0$ | 0.473 cm |
| Maximum muscle length / optimal muscle length | $\alpha_L$ | 1.1 |
| Minimum joint angle | $\theta_{min}$ | 70° |
| Maximum joint angle | $\theta_{max}$ | 140° |
| Parameters determine rotation range | $T_A^S$ | 1000 dyn |
| | $\sigma^S$ | 0.01 rad |
| Muscle activation time constants | $\tau_1$, $\tau_2$ | 0.01 s |
| Maximum muscle force | $F_0$ | $2.13 \times 10^5$ dyn |
| Normalization constant for motoneuron firing rate | $r_0$ | 54 Hz |
| Passive force | $c_1$ | 138 |
| | $k_1$ | 0.046 |
| | $L_{r1}$ | 1.17 |
| Length-dependent muscle activation | $\alpha_f$ | 0.56 |
| | $n_{f0}$ | 2.1 |
| | $n_{f1}$ | 3.3 |
| Active force-length relationship | $\beta_{FL}$ | 1.55 |
| | $\omega$ | 0.75 |
| | $\rho$ | 2.12 |
